# Testosterone acts through membrane protein GPRC6A to cause cardiac edema in zebrafish embryos

**DOI:** 10.1101/2023.03.20.533512

**Authors:** Vahid Zadmajid, Shayan Shahriar, Daniel A. Gorelick

## Abstract

Androgen actions are classically thought to be mediated by intracellular androgen receptors (AR), but they can also exert non-genomic effects via binding to integral membrane proteins. Although several putative membrane androgen receptors were cloned and characterized *in vitro*, their function as androgen receptors *in vivo* remains to be further investigated. Here, we used a chemical-genetic screen in zebrafish and found that the G-protein coupled receptor GPRC6A mediates non-genomic androgen action during embryonic development. Exposure to three androgens, 5α-Androstane-3,17-dione (androstanedione), dihydrotestosterone (DHT), and testosterone, caused cardiac edema or tail curvature in wild-type embryos. *ar* mutant embryos also exhibited cardiac edema or tail curvature following exposure to these androgens, suggesting the involvement of *ar*-independent pathways. To identify the causative receptor, we mutated putative membrane androgen receptors *gprc6a*, *hcar1-4*, or *zip9* genes and exposed mutant embryos to the androgens. We found that *hcar1-4* and *zip9* mutant embryos were susceptible to the identified androgens and developed cardiac edema or tail curvature phenotype following exposure. In contrast, we observed a significant reduction in cardiac edema phenotype in the *gprc6a* mutants compared to the wild-type embryos following testosterone treatment. Additionally, we exposed wild-type embryos to testosterone together with GPRC6A antagonists and observed a significant suppression of the cardiac edema phenotype. These results suggest that testosterone causes cardiac edema in zebrafish embryos by acting via the integral membrane protein GPRC6A, independently of nuclear androgen receptors. Using RNA-seq and RNA rescue approaches, we find that testosterone-GPRC6A causes cardiac phenotypes by reducing Pak1 signaling. Our study provides insights into non-genomic androgen signaling during embryonic development and identifies GPRC6A as a key receptor mediating androgen action.

## Introduction

Androgens regulate the development and function of multiple tissues and cell types, including muscle, heart, bone, skin, prostate, reproductive tract, and the brain (Bakker, 2022; Cunha et al., 2017; Murashima et al., 2015; Rossetti et al., 2017). Classically, androgens act by binding to cytosolic androgen receptors (AR), ligand-inducible transcription factors (Chang et al., 1988; Lubahn et al., 1988a; Lubahn et al., 1988b; Tilley et al., 1989; Trapman et al., 1988). Androgens can also activate signal transduction pathways independently of transcription, via a mechanism called non-genomic signaling (Gorczynska and Handelsman, 1995). These rapid androgen actions can be mediated by second messenger signaling pathways such as activation of intracellular kinases (eg MAPK, PI-3K) or increases in intracellular Ca^2+^ (Estrada et al., 2003; Gatson and Singh, 2007; Gatson et al., 2006; Guo et al., 2002; Kang et al., 2004; Lieberherr and Grosse, 1994; Liu et al., 2022). Androgen signaling occurs in the absence of nuclear AR (Guo et al., 2002; Hatzoglou et al., 2005; Pi et al., 2010). Such non-genomic androgen signaling has been observed in different cell types, including T cells and macrophages (Benten et al., 1999; Wunderlich et al., 2002), osteoblasts (Kang et al., 2004; Lieberherr and Grosse, 1994), prostate and breast cancer cells (Hatzoglou et al., 2005; Liu et al., 2015; Sun et al., 2006), granulosa and Sertoli cells (Gorczynska and Handelsman, 1995; Machelon et al., 1998), and skeletal muscle (Estrada et al., 2000). However, the receptors that mediate non-genomic androgen signaling are not well understood.

Several putative membrane androgen receptors such as G protein-coupled receptor family C group 6 member A (GPRC6A) (Pi and Quarles, 2012; Pi et al., 2008; Pi et al., 2010), the zinc transporter member 9 protein (ZIP9/SLC39A9) (Thomas et al., 2014), the G protein-coupled oxo-eicosanoid receptor 1 (OXER1) (Kalyvianaki et al., 2017), and ion channels such as transient receptor potential melastatin 8 (TRPM8) (Asuthkar et al., 2015a; Asuthkar et al., 2015b), or L-type Ca^2+^ channel, Cav1.2 (Scragg et al., 2004; Scragg et al., 2007) were identified based on their ability to bind androgens *in vitro*.

However, the evidence for androgen-dependent function of such proteins *in vivo* is less clear, in part because receptors have multiple ligands. For example, zebrafish *zip9* mutants exhibited reduced fecundity and egg viability compared with wild-type (Converse and Thomas, 2020) while mouse and zebrafish with mutations in *Cav1.2* exhibited cardiac abnormalities (Fu et al., 2011; Rosati et al., 2011; Rottbauer et al., 2001; Seisenberger et al., 2000), but whether these phenotypes are influenced by androgens, or other ligands binding to these receptors, is not known.

Here, we describe an *in vivo* screening approach combining chemical exposures and CRISPR/Cas9 genome editing in zebrafish embryos to discover membrane androgen receptors. Using this approach, we discovered that exposure to exogenous testosterone alters cardiac development in zebrafish embryos, and that this occurs via GPRC6A, independently of nuclear AR. We provide evidence that testosterone, and not a metabolite, acts via GPRC6A. We also discovered that exposure to two additional androgens, androstanedione and dihydrotestosterone, cause developmental phenotypes in zebrafish, independently of nuclear AR, GPRC6A, ZIP9 or OXER1, suggesting that additional membrane androgen receptors exist and influence embryonic development.

## Materials and Methods

### Zebrafish

Zebrafish were raised at 28.5°C on a 14-h light, 10-h dark cycle in the Baylor College of Medicine (BCM) Zebrafish Research Facility in a recirculating water system (Aquaneering, Inc., San Diego, CA) and a Tecniplast recirculating water system (Tecniplast USA, Inc., West Chester, PA). Wild-type zebrafish were AB strain (Westerfield, 2000) and all mutant lines were generated on the AB strain. The following previously published mutant and transgenic lines were used: nuclear androgen receptor *ar ^uab142^* (Crowder et al., 2018), estrogen receptor mutants *esr1a ^uab118^* (ERα), *esr2a ^uab134^* (ERβ1), *esr2b ^uab128^* (ERβ2) (Romano et al., 2017), and *Tg(myl7:EGFP)*(Huang et al., 2003). All procedures were approved by the BCM Institutional Animal Care and Use Committee.

### Embryo collection

Adult zebrafish were allowed to spawn naturally in pairs or in groups. Embryos were collected during 20 minute intervals to ensure precise developmental timing within a group. Embryos were placed in 60 cm^2^ Petri dishes at a density of no more than 100 per dish in E3B media (60X E3B: 17.2g NaCl, 0.76g KCl, 2.9g CaCl_2_-2H_2_O, 2.39g MgSO_4_ dissolved in 1 L Milli-Q water; diluted to 1X in 9 liter Milli-Q water plus 100 μL 0.02% methylene blue), and then stored in an incubator at 28.5°C. Between 24- and 48-hours post fertilization (hpf), embryos were manually dechorionated.

### Embryo treatments

At 2-4 hpf, embryos were incubated in E3B with androgen receptor modulator(s) or estradiol at 28.5°C until imaging or RNA extraction at 3 dpf. Dead embryos were removed daily. Embryos that were alive but abnormal were left in the dish and recorded. We did not observe any significant mortality differences between the treatments and vehicle control. All chemicals were purchased as follows: androstanedione CAS registry number 846-46-8, Isosciences catalog # 13386, purity ≥ 95%; dihydrotestosterone CAS 521-18-6, Isosciences catalog # 6065UNL, purity ≥ 98%; testosterone CAS 58-22-0, Sigma catalog # T1500, purity ≥ 98%; calindol CAS # 29610-18-8, Sigma catalog # SML1386, purity ≥ 98%; NPS-2143, CAS # 324523-20-8, Sigma catalog # SML0362, purity ≥ 98%; epigallocatechin gallate, CAS # 989-51-5, Sigma catalog # E4143, purity = 99.87%; estradiol, CAS # 50-28-2, Sigma catalog # E8875, purity ≥ 98%. All chemical stocks were made in 100% DMSO at 1000x and diluted in E3B embryo media to the final concentration (1x) at the time of treatment. All vehicle controls are 0.1% DMSO.

### CRISPR-Cas9 mutant generation

Chemically synthesized gRNAs for *gprc6a*, *zip9*, *oxer1* (*hcar1-4*), *cav1.2*, and purified Cas9 protein were obtained from Synthego (Redwood City, CA, USA). The target sequences were identified using CHOPCHOP (Labun et al., 2021) or zebrafish genome-scale lookup table from Wu et al. (Wu et al., 2018). Target site sequences are shown in Table S1. 1-cell-stage embryos were injected using glass needles pulled on a Sutter Instruments Fleming/Brown Micropipette Puller, model P-97, and a regulated air-pressure micro-injector (Harvard Apparatus, New York, PL1–90). Each embryo was injected with a 1 nl solution containing 1 µl of 10μM gRNA, 1 µl of 20μM Cas9 protein, 2 µl of 1.5 M KCl, 1 µl phenol red. The mixture was filled to 10 μL with 1X microinjection buffer (10mM Tris-HCL, 0.1mM EDTA, pH 8.0 in nuclease-free H_2_O) and incubated at 37 °C for 5-7 min before injection as described (Burger et al., 2016).

To generate *cyp19a1b* mutants, we hybridized oligonucleotides (oligo 1 5’-TAGGAGCATGTGGTAAAGGATG, oligo 2 5’-AAACCATCCTTTACCACATGCT) to generate double-stranded target DNA and annealed the DNA into digested pT7-gRNA plasmid. pT7-gRNA plasmid was digested simultaneously with BsmBI, BglII and SalI for one hour at 37 °C followed by one hour at 55 °C. gRNAs were synthesized and purified from this plasmid template using in vitro transcription (MegaShortScript T7 kit, Life Technologies). Cas9 mRNA was generated using in vitro transcription (mMessage mMachine T3 kit, Life Technologies) from a linearized pT3TS-nCas9n plasmid. RNA was purified using RNA clean & concentrator kit (Zymo Research). Plasmids pT7-gRNA and pT3TS-nCas9n were obtained from Addgene (numbers 46759, 46757)(Jao et al., 2013). Embryos were injected with a 1 nl solution containing 150 ng/μl of Cas9 mRNA, 50 ng/μl of gRNA and 0.1% phenol red.

For all gRNAs, mixtures were injected into the yolk of each embryo. Injected embryos were raised to adulthood and crossed to wild-type fish (AB strain) to generate F1 embryos. F1 offspring were sequenced via tail fin biopsies to identify mutations predicted to cause loss of functional protein.

### Genomic DNA isolation

Individual embryos or tail biopsies from individual adults were placed in 100 μL ELB (10 mM Tris pH 8.3, 50 mM KCl, 0.3% Tween 20) with 1 μL proteinase K (800 U/ml, NEB) in 96 well plates, one sample per well. Samples were incubated at 55°C for 2 hours (embryos) or 8 hours (tail clips) to extract genomic DNA. To inactivate Proteinase K, plates were incubated at 98°C for 10 minutes and stored at -20°C.

### High resolution melt curve analysis

PCR and melting curve analysis was performed as described (Parant et al., 2009). PCR reactions contained 1 μl of LC Green Plus Melting Dye (BioFire Diagnostics), 1 μl of Ex Taq Buffer, 0.8 μl of dNTP Mixture (2.5 mM each), 1 μl of each primer (5 μM), 0.05 μl of Ex Taq (Takara Bio Inc), 1 μl of genomic DNA, and water up to 10 μl. PCR was performed in a Bio-Rad CFX96 thermal cycler, using black/white 96 well plates (Bio-Rad HSP9665). PCR reaction protocol was 98°C for 1 min, then 34 cycles of 98°C for 10 sec, 60°C for 20 sec, and 72°C for 20 sec, followed by 72°C for 1 min. After the final step, the plate was heated to 95°C for 20 sec and then rapidly cooled to 4°C. Melting curves were generated with a Bio-Rad CFX96 Real-Time System over a 70–95°C range and analyzed with Bio-Rad CFX Manager 3.1 software. All mutations were confirmed by TA cloning and sequencing. PCR primer sequences are shown in Table S2.

### ELISA testosterone uptake assay

At 2-4 hpf, embryos were incubated in E3B with testosterone (30 µM) or vehicle controls (0.1% DMSO) at 28.5°C until 3 dpf. Embryos were collected at 3 dpf and transferred into 10 cryovials (Genesee Scientific, Cat #: 14-125) using a pipette (for each treatment 50 embryos were pooled and placed in each 1.5 mL cryovial). E3B media was aspirated out by pipetting (by immersing the pipette tips just below the liquid’s surface). Then, the embryos were washed 4 times using 1 ml 1X PBS, the media was aspirated out and the samples were frozen overnight at -80 ◦C. Embryos were thawed on ice and were homogenized in 200 µL of 1X PBS using a hand pellet pestle homogenizer (Genesee Scientific, Cat #: 27-118) and stored overnight at -20°C. After two additional freeze-thaw cycles to break the cell membranes (2 hours at -20°C and 10 minutes on ice), the homogenates were centrifuged for 10 minutes at 10000 x g, 4°C to collect the supernatant. The supernatant was dried in a vacuum evaporator (Thermo Scientific SpeedVac DNA130) and resuspended in 20 µL of 0.5% trifluoroacetic acid (TFA) in 5% acetonitrile (ACN) before processing with Pierce™ C18 Spin Columns (Thermo Scientific™, Cat #: 89870). Each column was activated with 200 µL of 50% ACN and centrifuged at 1,500 x g for 1 minute. The columns were equilibrated with 200 µL of 0.5% TFA, 5% ACN, and centrifuged at 1,500 x g for 1 minute. The full 20 µL samples were loaded on top of the resin bed in the column and centrifuged at 1,500 x g for 1 minute. The columns were washed by adding 200 µL of 0.5% TFA in 5% ACN and centrifuged at 1,500 x g for 1 minute. 20 µL of elution buffer (70% ACN) was added to the top of the resin bed and the columns were centrifuged at 1,500 x g for 1 minute. Finally, the samples were dried in a vacuum evaporator and resuspended in 200 µL of 1X PBS. Testosterone levels were quantified using ELISA kits according to the manufacturer (Cusabio Biotech, Cat #: CSB-E17554Fh). A serial dilution factor test was run for 1X, 100X, 500X, 1000X, and 2000X samples so that the concentration of testosterone in each sample was within the assay detection range (0.1 – 20 ng/mL). A standard curve was generated by measuring serially diluted standards provided by the manufacturer (S0 = 0 ng/mL; S1 = 0.1 ng/mL; S2 = 0.4 ng/mL; S3 = 1.6 ng/mL; S4 = 5 ng/mL; S5 = 20 ng/mL). Mean absorbance readings for each standard (in duplicate) were plotted on the x-axis against the concentration on the y-axis to draw a best-fit curve through the points on the graph using Excel (Harris Model: y=1/(a+bx^c); r = 0.99965). Samples were run in duplicate. Briefly, 50 μl of each sample, 50 μl of T-horseradish peroxidase conjugate (HRP-conjugate), and 50 μl of antibody were dispensed in each well of a 96-well plate and incubated for 1 hour at 37°C. After aspirating each well and 3 washes (400 μl per well using manufacturer’s 1X wash buffer), 50 μl of manufacturer’s substrate A and 50 μl of substrate B were added to each well. The plate was then incubated for 15 min at 37°C in the dark and the reaction was stopped by adding 50 μl of stop solution (1 N H_2_SO_4_). The optical density was read within 10 minutes at 450 nm with a microtiter plate reader (Cytation™ 5 Cell Imaging Multi-Mode Reader, Biotek, USA) and BioTek Gen5 Software. ELISA data of samples were interpolated from the standard curve generated by Excel to calculate the concentrations of target samples using the ELISA data analysis guide provided by CUSABIO (https://www.cusabio.com/m-225.html). Testosterone is presented as ng/ml. To calculate the amount of testosterone per embryo, we divided the total concentration of testosterone in the pooled embryos by 50. The detection limit of testosterone was 0.1 ng/ml, with sensitivity 0.05 ng/mL, according to the manufacturer.

### Live imaging

Embryos were imaged with a Nikon SMZ25 microscope equipped with a Hamamatsu ORCA-Flash4.0 digital CMOS camera, or with a Nikon SMZ18 microscope equipped with a Nikon DS-Fi3 camera. Images were equally adjusted for brightness and contrast in Adobe Photoshop Creative Cloud. For imaging of embryos following chemical exposure, embryos were imaged in E3B in 60 mm^2^ dishes in 0.04% tricaine anesthetic. For embryos in a mixed clutch containing multiple genotypes, embryos were genotyped following imaging using high resolution melting curve analysis. For confocal microscopy, embryos were anesthetized in E3B with 0.04% tricaine, positioned ventrally in 0.8% low-melt agarose in E3B on glass cover slips and imaged on a Nikon Ti2-E inverted microscope with Yokogawa CSU-W1 spinning disk confocal and Photometrics Prime95b sCMOS camera. Images were captured and maximum intensity projections were created using Nikon NIS-Elements-AR v5.42.03. Images were adjusted for brightness/contrast using FIJI (Schindelin et al., 2012).

### RNA Extraction and RNAseq Sample Preparation

Starting at 2-4 hpf, embryos were incubated in E3B with testosterone, testosterone + EGCG, testosterone + calindol and 0.1% DMSO (vehicle control) at 28.5°C until RNA extraction at 3 dpf. Dead embryos were removed daily. In both testosterone and testosterone + calindol (T+calindol) treated groups, embryos that exhibited cardiac phenotypes were pooled and selected for RNAseq. In the testosterone + EGCG (T+EGCG) treated groups, embryos that exhibited normal phenotypes were pooled and selected for RNAseq. All DMSO-treated embryos were normal. Treated embryos (20 per group) were transferred into RNAse-free tubes (Thermo-Scientific™ Snap Cap Low Retention Microcentrifuge Tubes, catalog #3434). Embryos were euthanized by placing the tube in ice and residual E3 media was withdrawn using 27 G x 1.25 inch BD PrecisionGlide™ Needle (catalog #305109). 200uL Trizol reagent (Thermo-scientific, catalog #15596026) was added to each tube containing treated embryos followed by homogenization with a motorized pestle. After that, an additional 800uL Trizol was added per tube, vortexed, and incubated at room temperature for 5 minutes. Tubes were centrifuged at 12000G for 10 minutes at 4°C. Then, we transferred the supernatant to fresh tubes and stored them at -80°C overnight. Frozen embryo lysates were thawed on ice the next day. We used the Direct-zol RNA miniprep Kit from Zymo Research (catalog #11-331) and followed the manufacturer protocol to extract RNA. RNA quality control analysis was performed using NanoDrop 1000 (Thermo-scientific) and Agilent 4200 TapeStation System. Good quality RNA with 260/280 absorbance of 2.0 or more and RIN value of 9.0 or more was sent for Directional mRNA library preparation (poly A enrichment) and NovaSeq X Plus Series (paired-end, 150 basepair max read length) sequencing by Novogene America.

### RNA Sequencing Analysis

Data analysis was performed on the high-performance computing (HPC) cluster that is managed by the Biostatistics and Informatics Shared Resource (BISR) at the Baylor College of Medicine. We trimmed the raw RNA sequencing FastQ data for low-quality reads using trim_galore (version 0.6.10) (Krueger, 2015), and then mapped them using STAR (version 2.7.1a) (Dobin et al., 2013) onto the zebrafish genome assembly GRCz11, maintained by the Genome Reference Consortium (GRC). Mapped sequence reads were aligned to genomic features using featureCounts (Liao et al., 2014). Combined genomic features were normalized using upper-quartile (UQ) and Remove Unwanted Variation method (Risso et al., 2014). Differentially expressed genes (DEGs) were determined using the EdgeR R package (Robinson et al., 2010). We made a separate list of DEGs (FDR < 0.05) for each treatment group (testosterone, testosterone + EGCG, testosterone + calindol) by comparing their genomic features with the genomic features of the control group treated with 0.1% DMSO. After that, we made a list of T-GPRC6A-specific genes (testosterone-GPRC6A-specific genes) by combining the significant genes that are differentially expressed in one direction (upregulated or downregulated in terms of fold change) in both testosterone and T+calindol treated groups but get rescued in the T+EGCG treated group by either showing no fold change or fold change in the opposite direction (e.g. if genes are upregulated in T, they will be downregulated in T+EGCG or vice versa) or 50% reduced fold change in the same direction (e.g. if up/downregulation (T) = 2x, up/downregulation (T+EGCG) ≤ x). GOBP (Gene Ontology Biological Process) pathway enrichment was performed using GSEA-v3.0 (https://www.gsea-msigdb.org/gsea/downloads.jsp) (Mootha et al., 2003; Subramanian et al., 2005). Any pathway with an adjusted FDR value <0.25 was considered significant. All RNA-seq data is deposited in the GEO repository, accession number GSE274852.

### Rescue experiment

CMV vectors (pTwist CMV) incorporating the open reading frame of either *pak1* or *grk3* were constructed by Twist Bioscience (San Francisco, CA, USA). All plasmids were sequenced prior to experimentation by Plasmidsaurus (Oregon, USA). Plasmids containing either *pak1* or *grk3* were linearized by the restriction enzyme NotI (NEB R3189S). mRNA was transcribed using the mMessage mMachine SP6 Kit (Invitrogen, AM1340) and purified with the RNA clean & concentration kit (Zymo Research, R1019). Embryos were microinjected at the one-cell stage with 1 nl total volume containing either *pak1* or *grk3* (30 pg mRNA) or *GFP* mRNA (50 pg mRNA). All solutions were injected into the yolk.

### Experimental design and data analysis

Each experiment included comparing groups (treated vs untreated or mutant vs wildtype) using 11-160 embryos per group with all embryos from the same clutch. Experiments were performed on 3-6 clutches for each genotype. This is essentially a complete block design with clutch as block. Mean cardiac edema or tail curvature rate of individual embryos from a clutch was used for comparing treatment groups (or mutant groups) within experiments using one-way ANOVA with Tukey’s multiple comparisons test. For some individual pairs of comparisons, paired t test was used. Significance level is 0.05. All Statistical analyses were performed using Prism version 9.4.1 (GraphPad Software, Boston, MA).

## Results

### Strategy to identify membrane androgen receptors using genetic and pharmacologic approaches

The strategy we adopted is shown schematically in Fig. 1. We exposed wild-type embryos to multiple androgens at different concentrations (10-100 μM) in embryo media, beginning at 3 hours post fertilization. We assayed embryo morphology at 2- and 3-days post fertilization (see Materials and Methods). We identified three androgens, 5α-Androstane-3,17-dione (androstanedione) (15 μM), dihydrotestosterone (DHT) (30 μM), and testosterone (30 μM), that caused a morphologic abnormality. All three compounds caused cardiac edema with elongated and incompletely looped hearts (Figs 2 and S1, Table 1; androstanedione mean embryos with cardiac edema 84% ± 12% standard deviation, DHT 96% ± 6%, testosterone 79% ± 4%, n=6 clutches, 15-27 embryos per clutch). Additionally, androstanedione and DHT, but not testosterone, caused a tail curvature phenotype in which the caudal tail was curved dorsally (androstanedione mean embryos with tail curvature 94% ± 5% standard deviation, DHT 81% ± 20). For embryos exposed to androstanedione or DHT, every embryo that exhibited cardiac edema also exhibited a tail curvature phenotype (Table 1). Other features of embryos, like gross morphology of the head, eyes, somites, otolith, notochord, brain ventricles, fin folds, and yolk, appeared normal, suggesting that these androgens affect specific developmental processes and do not cause generalized, non-specific toxicity.

**Figure 1.**
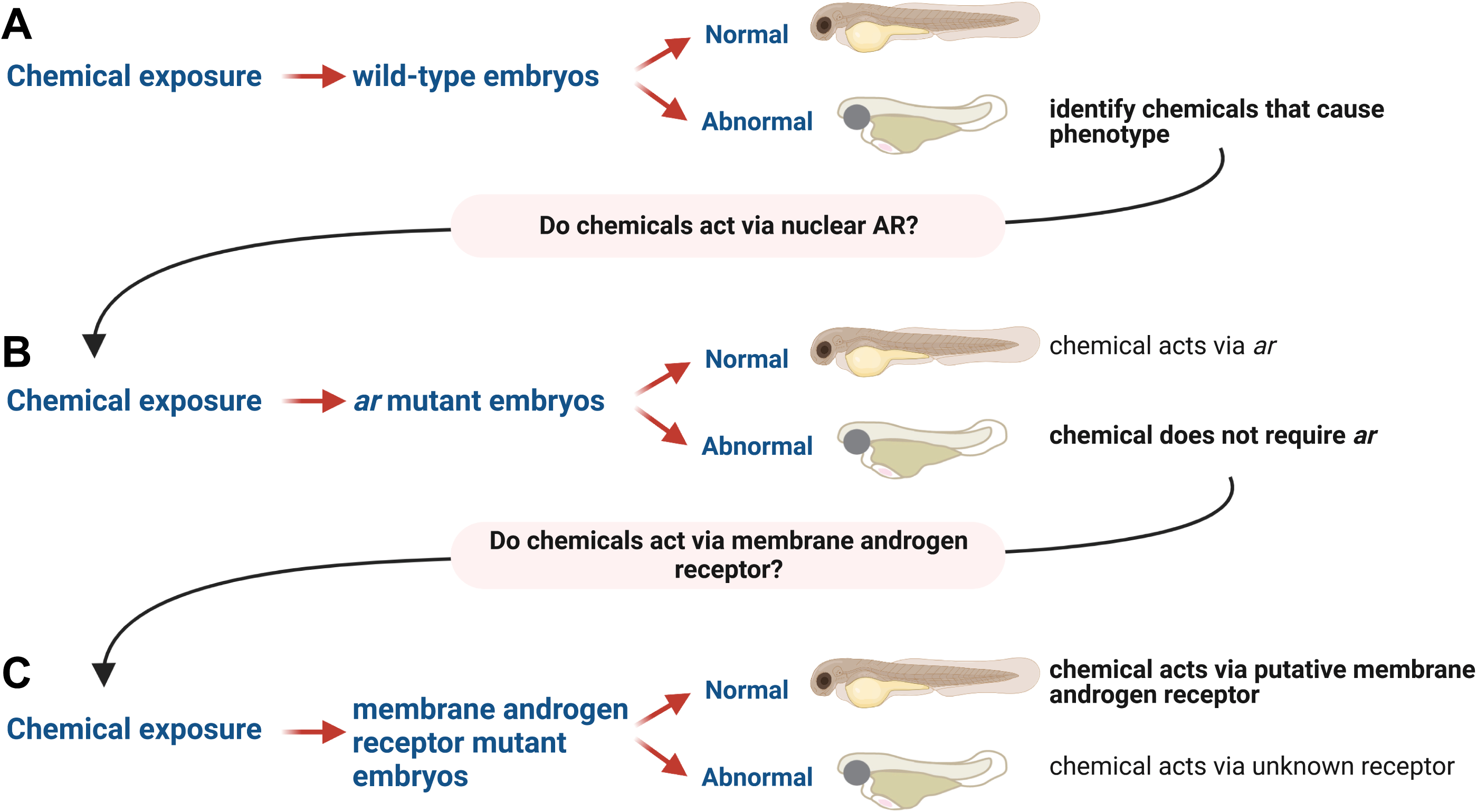
Chemical-genetic screen to identify receptors that mediate androgen-dependent phenotypes. **(A)** We exposed wild-type embryos to multiple androgens at different concentrations beginning at 3 hours post fertilization. We assayed embryo morphology at 2- and 3-days post fertilization. We identified chemicals that caused morphologic abnormalities. (B) We tested whether the phenotype is rescued in zebrafish with mutations in the nuclear androgen receptor gene (ar). If the phenotype is rescued, then the steroid acts via AR. **(C)** If the phenotype persists in the nuclear *ar* mutant, then we test zebrafish with mutations in membrane steroid receptors. To efficiently generate membrane androgen receptor mutants, we inject embryos with Cas9 and guide RNAs, expose injected embryos to chemicals, and screen for phenotypes. Following phenotypic screening, embryos are genotyped to determine mutation burden. Figure created using BioRender.com.

**Figure 2.**
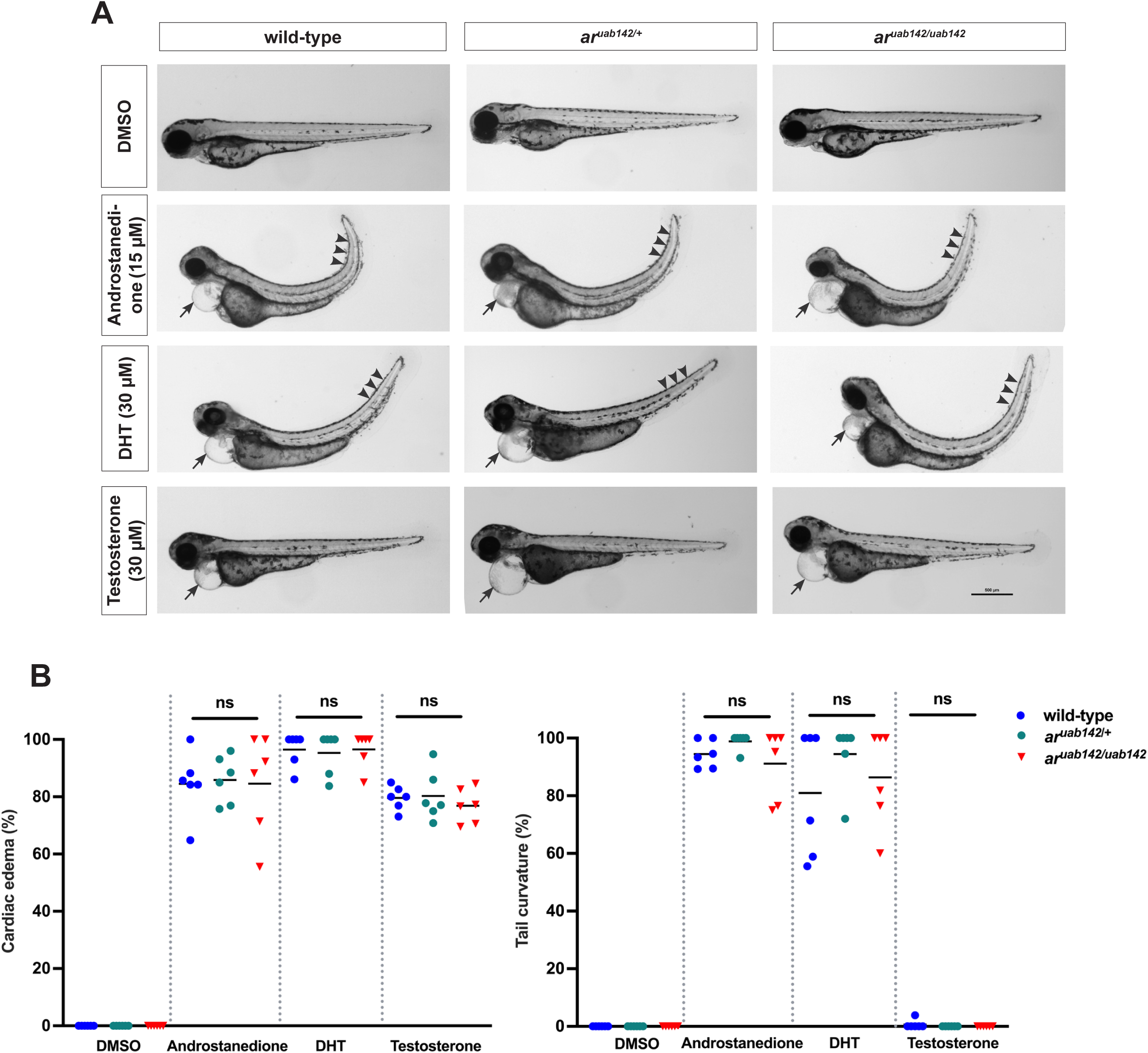
Identification of androgens that cause phenotypes in wild-type and nuclear androgen receptor (*ar*) mutant embryos. Wild-type or ar mutant embryos were exposed to vehicle (0.1% DMSO), 5α-Androstane-3,17-dione (androstanedione), dihydrotestosterone (DHT), or testosterone beginning at 3 hours post fertilization. (A) Representative images of embryos at 3 days post fertilization. Note cardiac edema (arrows) and tail curvature (arrowheads). Each clutch was derived from heterozygous parents and contained a mix of wild-type, heterozygous ar uab142/+, or homozygous ar uab142/uab142 embryos. Lateral views with anterior to the left, dorsal to the top. Scale bar = 500 μm. (B) Percent of embryos exhibiting cardiac edema or tail curvature. Each data point is the mean percent of embryos in a single clutch exhibiting cardiac edema or tail curvature (11-57 embryos per clutch). Clutches in the same treatment group were assayed on different days. Genotypes and treatments are separated by vertical gray dotted lines in each graph. Horizontal lines are the mean of each treatment. ns, not significant (p > .05), one-way ANOVA with Tukey’s multiple comparisons test.

**Table 1.**
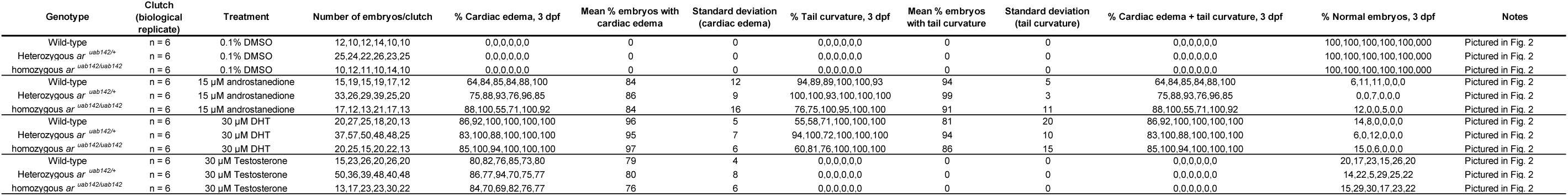
Embryo data accompanying Figure 2.

### Neither androstanedione, DHT, nor testosterone act via nuclear androgen receptor (*ar*) to cause morphologic phenotypes in zebrafish embryos

To explore whether these three androgens act via nuclear *ar* to cause morphologic phenotypes, we crossed heterozygous *ar ^uab142/+^* zebrafish to each other and exposed the resulting embryos to vehicle (0.1% DMSO), androstanedione, DHT, or testosterone beginning at 3 hours post fertilization. Following chemical exposure, we observed no statistically significant difference in the percent of embryos exhibiting cardiac edema or tail curvature phenotypes between wild-type, heterozygous *ar ^uab142/+^*, or homozygous *ar ^uab142/uab142^*embryos (Fig. 2, Table 1). We conclude that zebrafish embryos with mutations in nuclear *ar* are sensitive to androstanedione, DHT, or testosterone exposure, which raises the possibility that these androgens act via an *ar*-independent pathway to cause morphologic phenotypes during embryonic development.

### Quantification of testosterone uptake in zebrafish

Our androgen exposure concentrations in fish water (30 μM) is higher than the dissociation constant (Kd) for testosterone binding to the zebrafish nuclear *ar* (2 nM)(Jørgensen et al., 2007). However, it is not known how much of the androgen in the water is absorbed and bioavailable in the zebrafish embryo. We used ELISA to measure the amount of testosterone in zebrafish following exposure to water containing 30 μM testosterone from 0 to 3 dpf. We found that the testosterone concentration in the whole embryo was 65.90 ± 33.93 nM/per embryo (mean ± SD, Fig. 3). While this is ∼30 fold more than Kd for testosterone binding to the zebrafish AR/NR3C4 (2 nM), these levels are more similar to physiologic levels of circulating testosterone in adult human men (10-30 nM) (Burger, 2002; Finkelstein et al., 2013; Kelsey et al., 2014; Mohr et al., 2005; Travison et al., 2017; Wu et al., 2010). Moreover, this estimation does not consider the percent of androgens that are bioavailable or bound to serum globulins, which is not known in zebrafish embryos (Miguel-Queralt et al., 2004).

**Figure 3.**
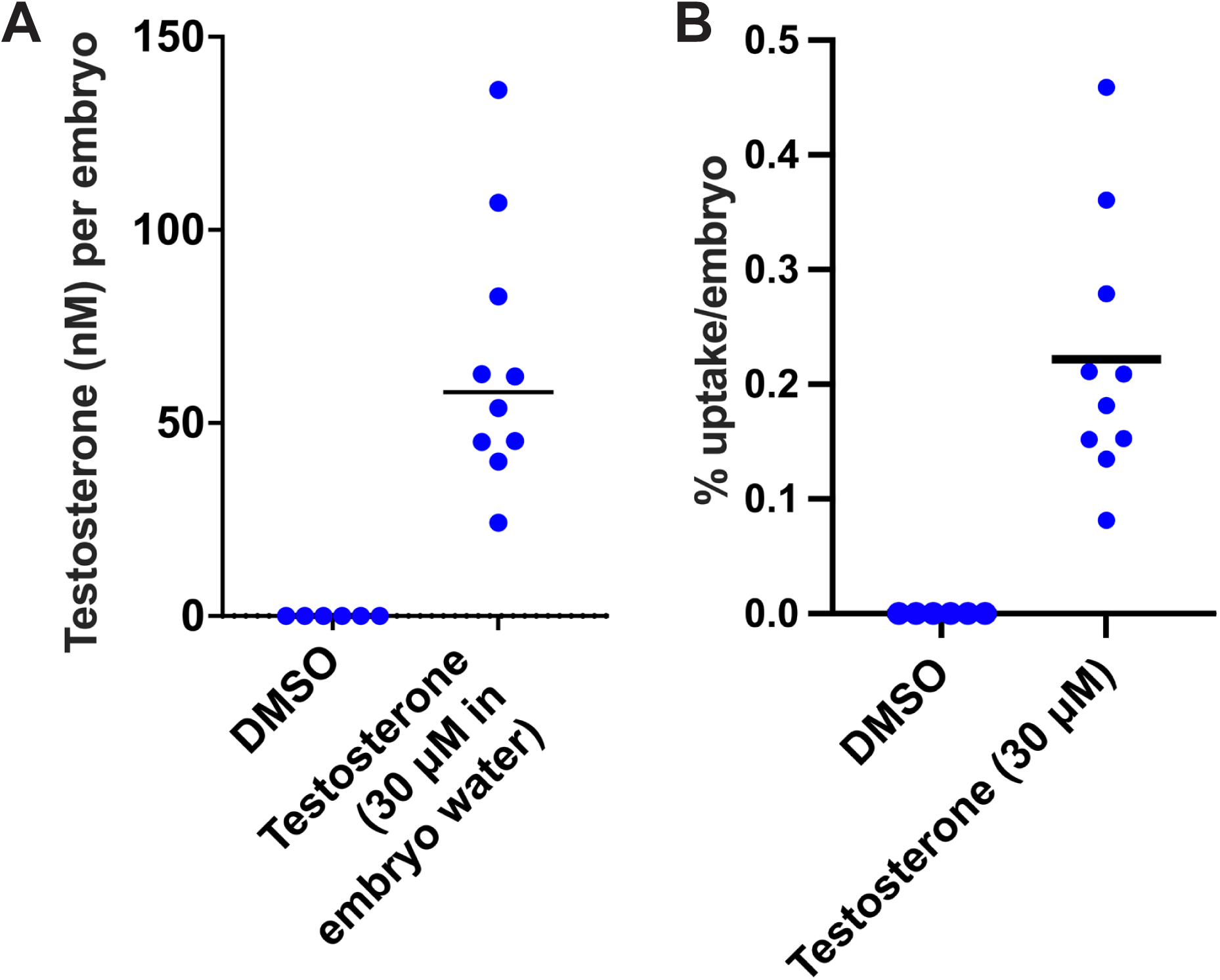
Testosteone concentration and uptake in zebrafish embryos. Wild-type embryos at 3 hours post fertilization (hpf) were exposed to vehicle (0.1% DMSO) or testosterone (30 μM) until 3 days post fertilization (dpf). Testosterone levels were measured using ELISA to calculate nM testosterone per embryo (A) and percent testosterone uptake per embryo (B). Each circle represents the mean testosterone concentrations or uptake per embryo for a single clutch of embryos, with 50 embryos pooled per clutch). Horizontal lines are the mean of each clutch.

We also calculated the percent uptake of testosterone per embryo. We found that the percent uptake was 0.22% ± 0.11% per embryo (mean ± SD, Fig. 3). The low uptake from water is consistent with our previous results for the structurally similar steroid, 17β-estradiol, where we found that 1% or less of estradiol in fish water is taken up into embryos, depending on embryo stage (Souder and Gorelick, 2017). Note that we were unable to detect testosterone in untreated embryos. We conclude that endogenous testosterone levels at these stages of embryonic development are below the limit of detection, at least when pooling 50 embryos. Taken together, our results suggest that only a small percentage of testosterone in the water is absorbed by the embryo, and that supra-physiologic concentrations of testosterone in the fish water are justified because it results in close to human physiologic levels in embryos.

### Testosterone, but not androstanedione or DHT, act via GPRC6A to cause cardiac edema

In the experiments described above, we showed that nuclear *ar* mutant embryos remained sensitive to androstanedione, DHT, or testosterone exposure, suggesting the involvement of *ar*-independent pathways. To determine whether these androgens act via putative membrane androgen receptors, we took an F0 CRISPR genome editing approach to generate zebrafish with mosaic mutations in putative membrane androgen receptor genes: *gprc6a*, *zip9*, *oxer1* (*hcar1-4), cacna1c* (*cav1.2*) (Thomas, 2019; Treviño and Gorelick, 2021)(Fig. S2). This approach enables us to inject embryos with a cocktail of 4 guide RNAs together with Cas9 protein, targeting the same gene redundantly, which induces mutations at a high enough burden to observe phenotypes in the injected embryos (Hoshijima et al., 2019; Kroll et al., 2021; Wu et al., 2018).

We exposed embryos injected with gRNAs targeting *gprc6a*, *zip9*, *hcar1-4, cacna1c*, or control embryos injected with Cas9 alone, to androstanedione, DHT, testosterone or vehicle (0.1% DMSO) beginning at 3 hours post fertilization and assayed morphology at 2- and 3-days post fertilization. Following androstanedione, DHT, or testosterone exposure, *hcar1-4* and *zip9* mutant embryos exhibited cardiac edema or tail curvature phenotype (Fig. 4). Untreated *cacna1c* mosaic mutant embryos exhibited cardiac edema with abnormal heart morphology at 3 days post fertilization (not shown). Therefore, we excluded *cacna1c* from further screening. Following chemical exposure, we found no statistically significant difference in the proportion of embryos with cardiac edema or tail curvature phenotype between control or *hcar1-4* and *zip9* F0 mosaic mutants (Fig 4, Table 2).

**Figure 4.**
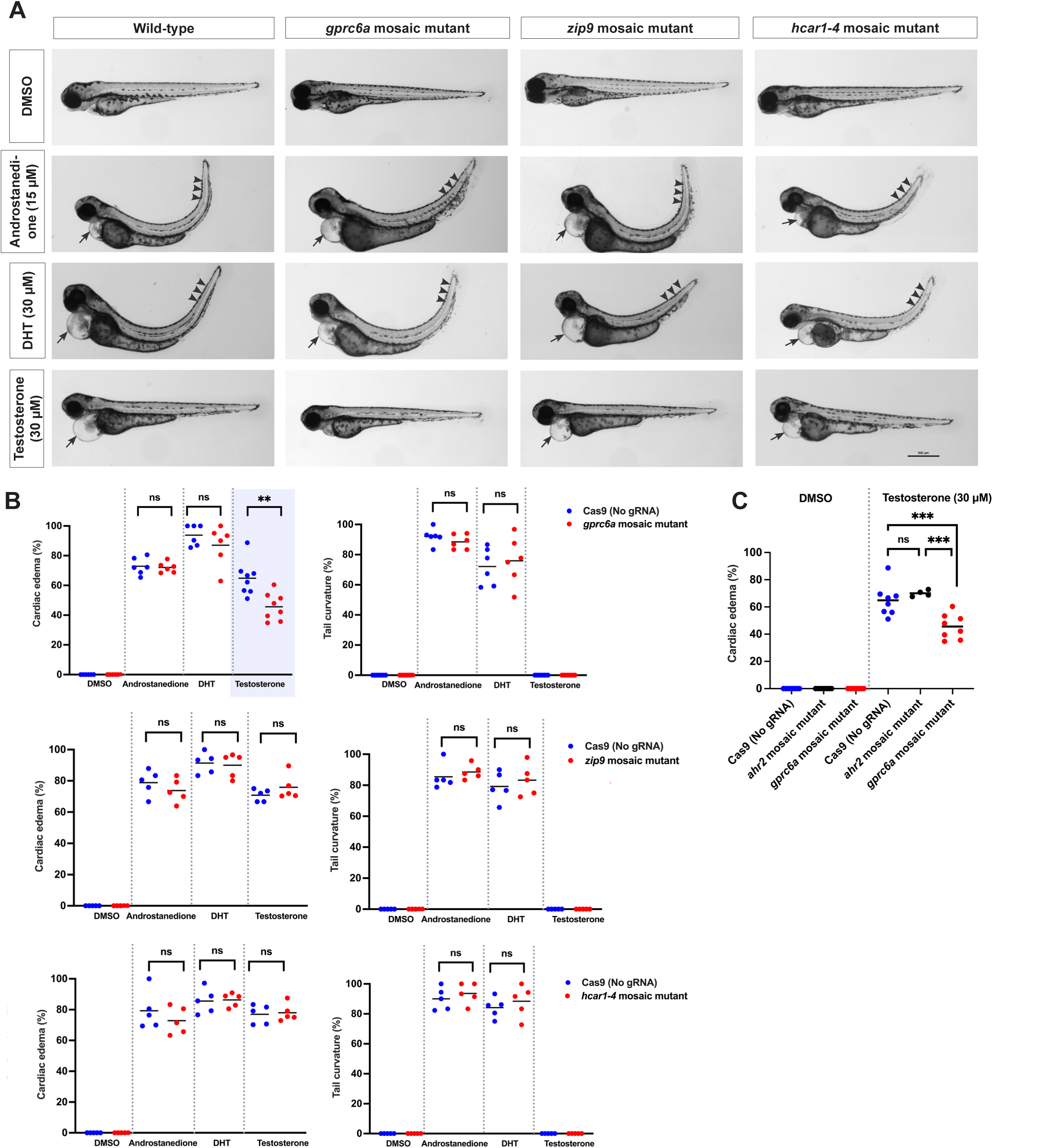
Testosterone phenotype is partially rescued by CRISPR targeting *gprc6a*. 1-cell stage embryos were injected with Cas9 protein alone or together with a cocktail of 4 guide RNAs targeting different regions of the indicated genes (*gprc6a*, *zip9*, *hcar1-4*). At 3 hours post fertilization, embryos were exposed to vehicle (0.1% DMSO), 5α-Androstane-3,17-dione (androstanedione), dihydrotestosterone (DHT), or testosterone. **(A)** Representative images of embryos at 3 days post fertilization. Note cardiac edema (arrows) and curved tail (arrowheads), except in *gprc6a* mosaic mutants following testosterone exposure. Lateral views with anterior to the left, dorsal to the top. Scale bars = 500 μm. **(B)** Percent of embryos exhibiting cardiac edema (left graphs) or tail curvature (right graphs). Each data point is the percent of embryos in a single clutch exhibiting cardiac edema or tail curvature (25-160 embryos per clutch). Clutches in the same treatment group were assayed on different days. Genotypes and treatments are separated by vertical gray dotted lines in each graph. Horizontal lines are the mean of each treatment. **(C)** As an additional control, embryos were injected with Cas9 protein alone or with guide RNAs targeting an unrelated gene *(ahr2),* and then exposed to vehicle or testosterone. The percent of embryos with cardiac edema was assayed at 3 days post fertilization. Each data point represents a single clutch (35-70 embryos per

**Table 2.**
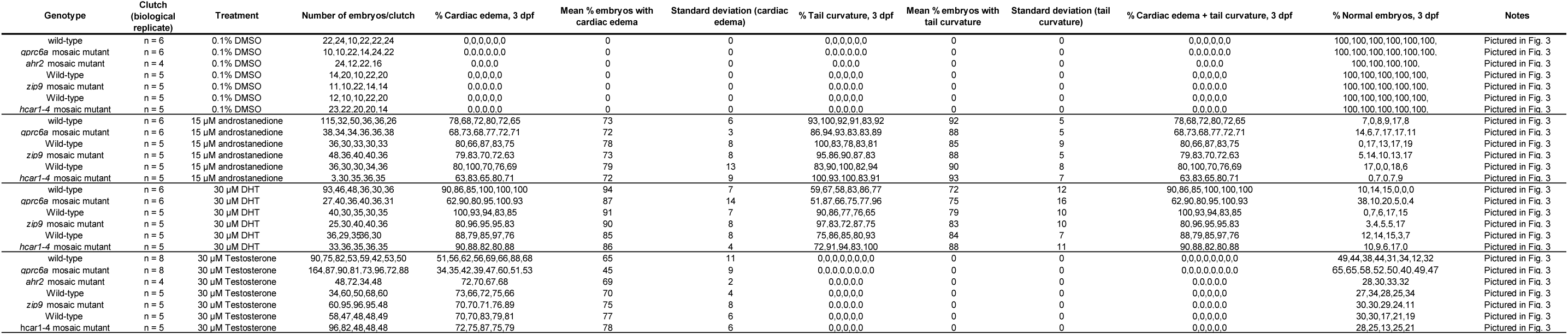
Embryo data accompanying Figure 4. Wild-type embryos were injected with Cas9 protein without any gRNAs

We found that *gprc6a* mosaic mutants had a statistically significant reduction in the percent of embryos exhibiting cardiac edema following testosterone exposure (Fig. 4B, Table 3; embryos injected with Cas9 alone mean cardiac edema 65% ± 11%, *gprc6a* mosaic mutants 45% ± 9%, n=8 clutches, 42-164 embryos per clutch). However, the *gprc6a* mosaic mutants remained sensitive to androstanedione or DHT exposure (Fig. 4B, Table 2). As an additional control, we injected wild-type embryos with Cas9 plus gRNAs targeting an unrelated gene encoding an aryl hydrocarbon receptor *(ahr2)*.

**Table 3.**
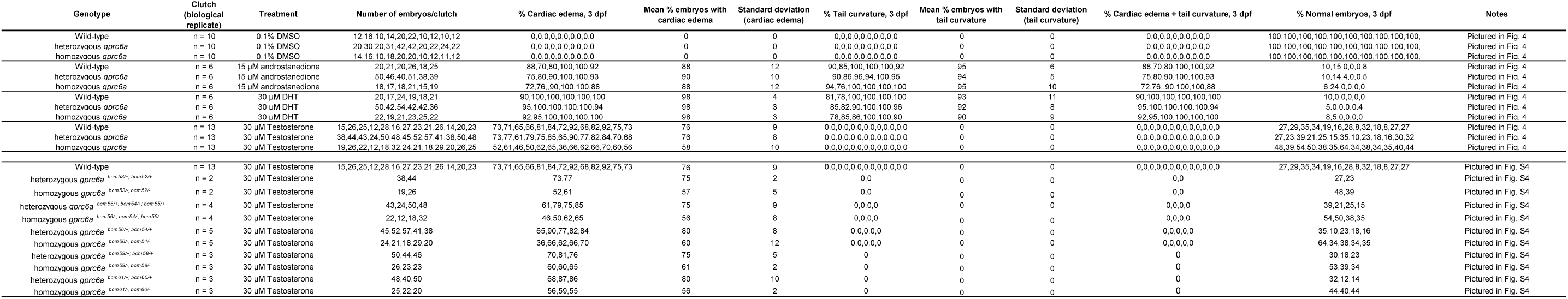
Embryo data accompanying Figure 6 and Figure S5. The same data are pooled and grouped by gene (top table) or not pooled and grouped by allele (bottom table)

We exposed *ahr2* mosaic mutant embryos to testosterone and assayed cardiac edema phenotype at 3 days post fertilization. There was no statistically significant difference in the percent embryos exhibiting cardiac edema between control embryos versus *ahr2* mosaic mutant embryos (Fig. 4C, Table 2). We conclude that the testosterone exposure phenotype is partially rescued by mutating *gprc6a*.

To test whether there is crosstalk between GPRC6A and the canonical nuclear androgen receptor *ar* (also known as *nr3c4*), we generated mosaic double mutant embryos by injecting 1-cell stage *ar^uab142/uab142^*embryos with a cocktail of 4 guide RNAs targeting the *gprc6a* gene (the same gRNAs used above). We exposed double mutant embryos to testosterone (30 μM) to test whether the phenotype is the same as in single mutant embryos. There was no statistically significant difference in the percent embryos exhibiting cardiac edema between *gprc6a* mosaic mutants on wild-type background versus *gprc6a* mosaic mutants on *ar^uab142/uab142^* mutant background (Fig. 5). We conclude that testosterone acts through *gprc6a* independently of nuclear *ar.* This is consistent with results from cultured cells and adult mouse tissues suggesting that GPRC6A responds to testosterone (Pi et al., 2010; Pi et al., 2015).

**Figure 5.**
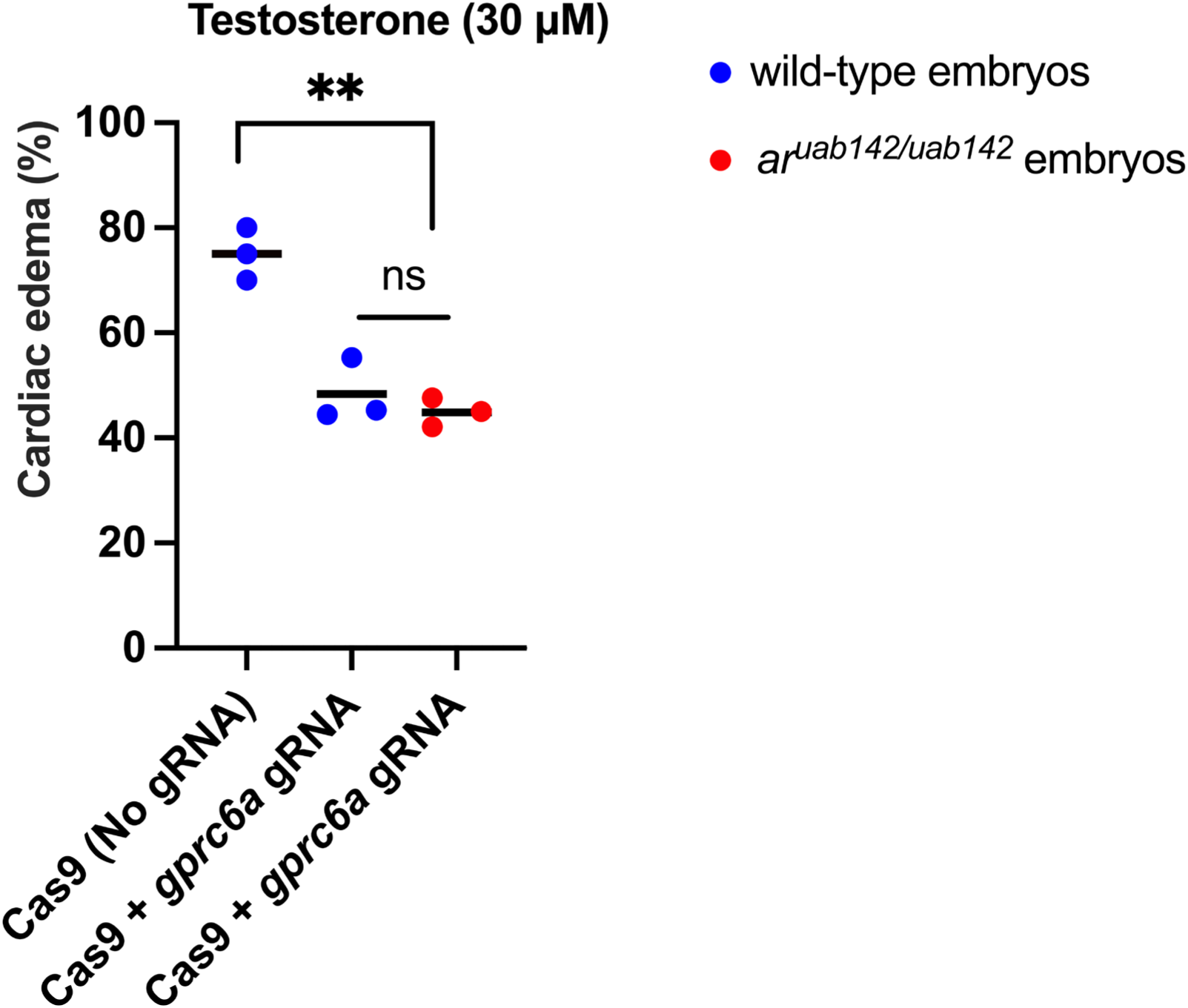
Testosterone phenotype in double mutants (*gprc6a* mosaic mutant + *ar^uab142/uab142^*) is similar to *gprc6a* mosaic mutant. 1-cell stage *ar^uab142/uab142^* embryos were injected with Cas9 protein alone or together with a cocktail of 4 guide RNAs targeting different regions of the *gprc6a* gene. At 3 hours post fertilization, embryos were exposed to testosterone. Morphologic phenotypes were assayed at 3 days post fertilization. Each data point is the mean percent of embryos in a single clutch exhibiting cardiac edema (19-80 embryos per clutch). Clutches in the same treatment group were assayed on different days. Horizontal lines are the mean of each group. **p < .01, ns, not significant (p > .05), one-way ANOVA with Tukey’s multiple comparisons test.

### CRISPR–Cas9 mutagenesis efficiency in F0 embryos

The absence of a phenotype in a CRISPR screen could be due to low efficiency of gRNAs. It is possible that targeting *zip9* or *hcar1-4* failed to rescue androgen phenotypes because the *zip9* and *hcar1-4* gRNAs had low efficacy. To evaluate the mutagenesis efficiency in our F0 embryos, we isolated genomic DNA from individual embryos at 24 hours post fertilization. The targeted site was amplified using PCR. Amplicons were Sanger sequenced and then deconvoluted and analyzed using the ICE (Inference of CRISPR Edits) algorithm (Conant et al., 2022). We compared indel mutations (the percent of sequence from each embryo that has a mutation) or knockout score (the percent of mutations from each embryo that were predicted to produce a non-functional protein) for each gRNA (Fig. S3). Different guides acted with variable effectiveness. In the cocktail of 4 guide RNA sets to target *gprc6a*, G4 produced an average of more than 50% indel mutations or knockout score (n=38 embryos from 2 clutches). G1 and G2 were less effective, with an average mutation frequency of less than 10% (n=40 embryos from 2 clutches), and G3 produced 0% mutation (n=38 embryos from 2 clutches) (Fig. S3).

Comparing the efficiency of guide RNAs targeting *zip9*, G1, and G3 were the most effective. G1 induced an average of 78% indel mutations (n=40 embryos from 2 clutches) or 52% knockout score (n=40 embryos from 2 clutches), and G3 induced an average of 73% indel mutations (n=45 embryos from 2 clutches) or 5% knockout score (n=45 embryos from 2 clutches). For *hcar1-4*, the average indel mutations or knockout score for all G1, G2, and G4 was greater than 50% (n=2 clutches/24-45 embryos per clutch). In analyzed *hcar1-4* mosaic mutant embryos, G4 induced greater than 90% indel mutations or knockout-score (n=46 embryos from 2 clutches). Overall, these results confirm the efficacy of multi-locus targeting to generate zebrafish F0 mutants for the purpose of initial screening of gene function.

### Zygotic *gprc6a* mutants recapitulate phenotypes observed in mosaic mutants

To validate our observations, we generated heritable, non-mosaic mutations in *gprc6a* and determined whether homozygous embryos were protected from testosterone-dependent cardiac edema, as were F0 mosaic mutants. We injected wild-type embryos with Cas9 together with the same set of 4 guide RNAs, targeting different coding regions of the *gprc6a* gene, as we did to generate *gprc6a* mosaic mutants (Fig. S4). Injected embryos were raised to adulthood and outcrossed to wild-type zebrafish to generate heterozygous F1 embryos. Heterozygous F1 adult fish were sequenced to identify mutations predicted to cause loss of functional GPRC6A protein. We identified several mutant alleles in F1 fish and derived five independent *gprc6a* mutant lines, from which we performed experiments on homozygous embryos: *gprc6a ^bcm53/-; bcm52/-^*, *gprc6a bcm56/-; bcm54/-; bcm55/-*_, *gprc6a*_ *bcm56/-; bcm54/-*_, *gprc6a*_ *bcm59/-; bcm58/-*_, and *gprc6a*_ *bcm61/-; bcm60/-*(Fig. S4A). For example, *bcm53* is a 1 bp deletion in exon 1 resulting in a frameshift at amino acid 44 and a premature stop codon at amino acid 50 in the ligand binding domain. *bcm5*2 is a 1 bp deletion in exon 3 resulting in a frameshift at amino acid 225 and a premature stop codon at amino acid 250. These two alleles are genetically linked (we were unable to generate fish containing *bcm52* without *bcm53*), however the upstream allele (*bcm53*) is likely driving the loss of function. We did not observe any morphologic differences among untreated *gprc6a ^bcm53/-; bcm52/-^*, *gprc6a ^bcm56/-; bcm54/-;^ bcm55/-*_, *gprc6a*_ *bcm56/-; bcm54/-*_, *gprc6a*_ *bcm59/-; bcm58/-*_, and *gprc6a*_ *bcm61/-; bcm60/-* _embryos._

We exposed homozygous *gprc6a* embryos to vehicle, androstanedione, DHT, or testosterone beginning at 3 hours post fertilization. Each clutch contained a mix of wild-type and mutant embryos (denoted as wild-type +/+, heterozygous +/-, or homozygous - /-; Fig. 6A). Following testosterone exposure, homozygous *gprc6a* mutant embryos exhibited partial cardiac edema rescue, similar to what we observed in mosaic F0 mutants (Figure 4B, Table 3; wild-type mean percent cardiac edema 76% ± 9%, heterozygous 76% ± 8%, homozygous 58% ± 10%, n=13 clutches, 12-57 embryos per clutch). Analyzing the response of each allele separately revealed similar results (Fig. S5, Table 3). Homozygous *gprc6a* mutants remained sensitive to androstanedione and DHT exposure, similar to wild-type embryos (Fig. 6B, Table 3). We conclude that the F0 CRISPR *gprc6a* mosaic mutants recapitulate what occurs in non-mosaic homozygous mutants. Taken together, our results suggest that testosterone causes cardiac edema by acting via the integral membrane protein GPRC6A independently of the nuclear *ar*.

**Figure 6.**
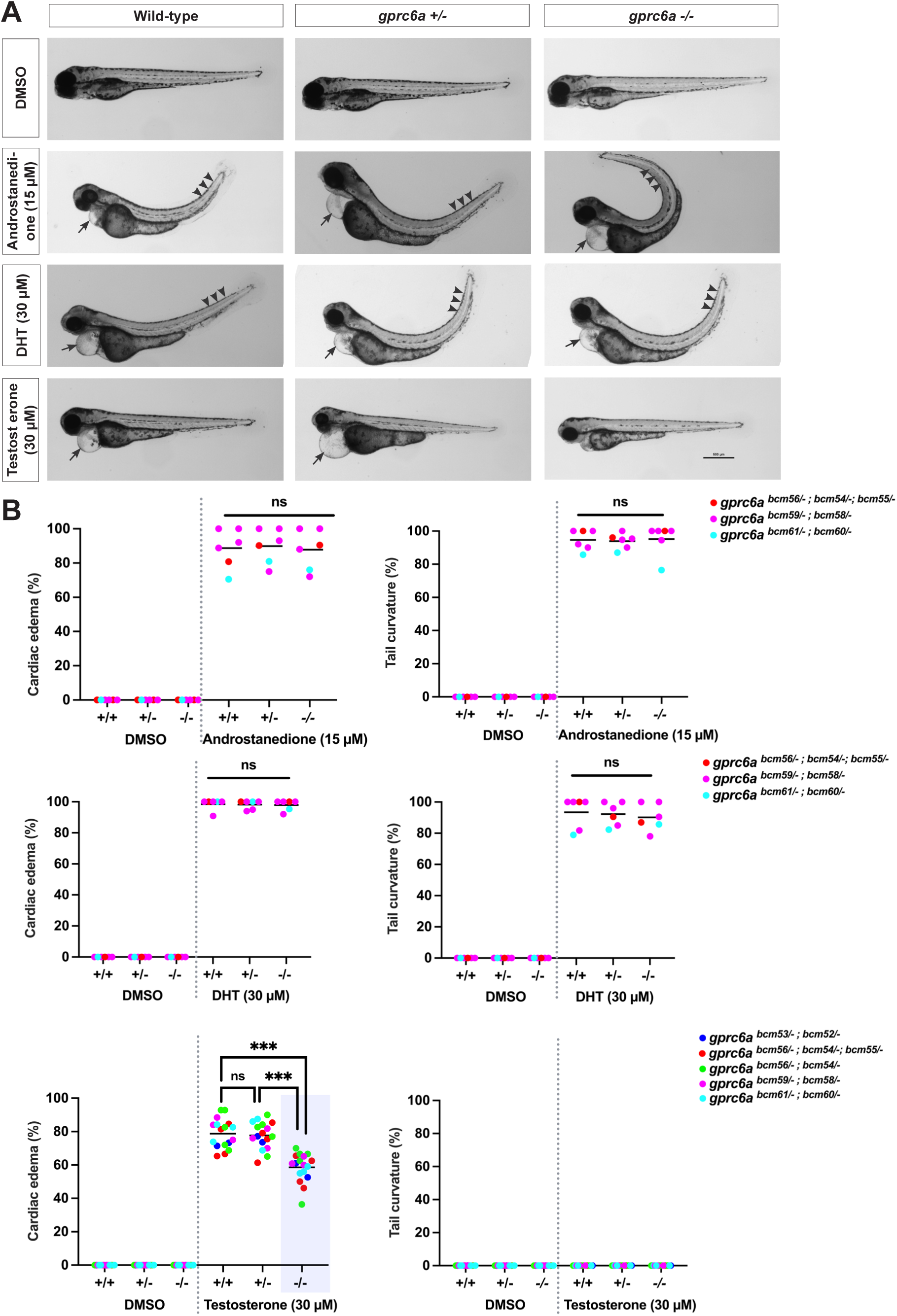
Testosterone phenotype is partially rescued in homozygous *gprc6a* mutants. Heterozygous *gprc6a* adult zebrafish were crossed to each other to generate zygotic *gprc6a* mutants. The clutch contained a mix of wild-type and mutant embryos (denoted as wild-type +/+, heterozygous +/-, or homozygous -/-). Embryos were exposed to vehicle (0.1% DMSO) or 5α-Androstane-3,17-dione (androstanedione), dihydrotestosterone (DHT), and testosterone beginning at 3 hours post fertilization. Morphologic phenotypes were assayed at 3 days post fertilization, then each embryo was genotyped. **(A)** Representative images of *gprc6a ^bcm56/-; bcm54/-; bcm55/-^* embryos at 3 days post fertilization. Note cardiac edema (arrows) and curved tail (arrowheads), except in homozygous *gprc6a* mutants following testosterone exposure. Lateral views with anterior to the left, dorsal to the top. Scale bars = 500 μm. **(B)** Percent of embryos exhibiting cardiac edema (left graph) or tail curvature (right graph). Each data point is the mean percent of embryos in a single clutch exhibiting cardiac edema or tail curvature (15-50 embryos per clutch). Clutches in the same treatment group were assayed on different days.

### GPRC6A antagonists inhibit testosterone-dependent cardiac edema in wild-type embryos

To further confirm that testosterone acts through *gprc6a* to cause cardiac edema, we tested whether GPRC6A antagonists block testosterone-driven cardiac edema. We exposed wild-type embryos to testosterone together with different GPRC6A antagonists, NPS-2143 (5 µM), calindol (5 µM), and epigallocatechin gallate (EGCG, 100 µM) at concentrations used to study GPRC6A responses in cultured cells (Faure et al., 2009; Mizuta et al., 2021; Pi et al., 2018). We confirmed that 5 µM NPS-2143, 5 µM calindol, and 100 µM EGCG do not cause abnormalities in embryos in the absence of androgen exposure (Fig. 7A, Table 4). We found that wild-type embryos exposed to NPS-2143 or EGCG with testosterone had statistically significantly reduced cardiac edema compared to embryos exposed to testosterone alone (Fig. 7A, Table 4; testosterone mean embryos with cardiac edema 69% ± 7%, testosterone + NPS-2143 39% ± 9%, testosterone + EGCG 32% ± 8%, n=6 clutches, 43-63 embryos per clutch). While NPS-2143 and EGCG partially blocked testosterone-driven cardiac edema, calindol failed to reduce testosterone-dependent cardiac edema (Fig. 7A, Table 4). Calindol was developed as a calcimimetic that binds the human calcium sensing receptor and has lower selectivity towards GPRC6A (Gloriam et al., 2011; Kessler et al., 2004). The failure of calindol to inhibit zebrafish GPRC6A could be due to the lower selectivity together with differences between zebrafish and human GPRC6A proteins.

**Figure 7.**
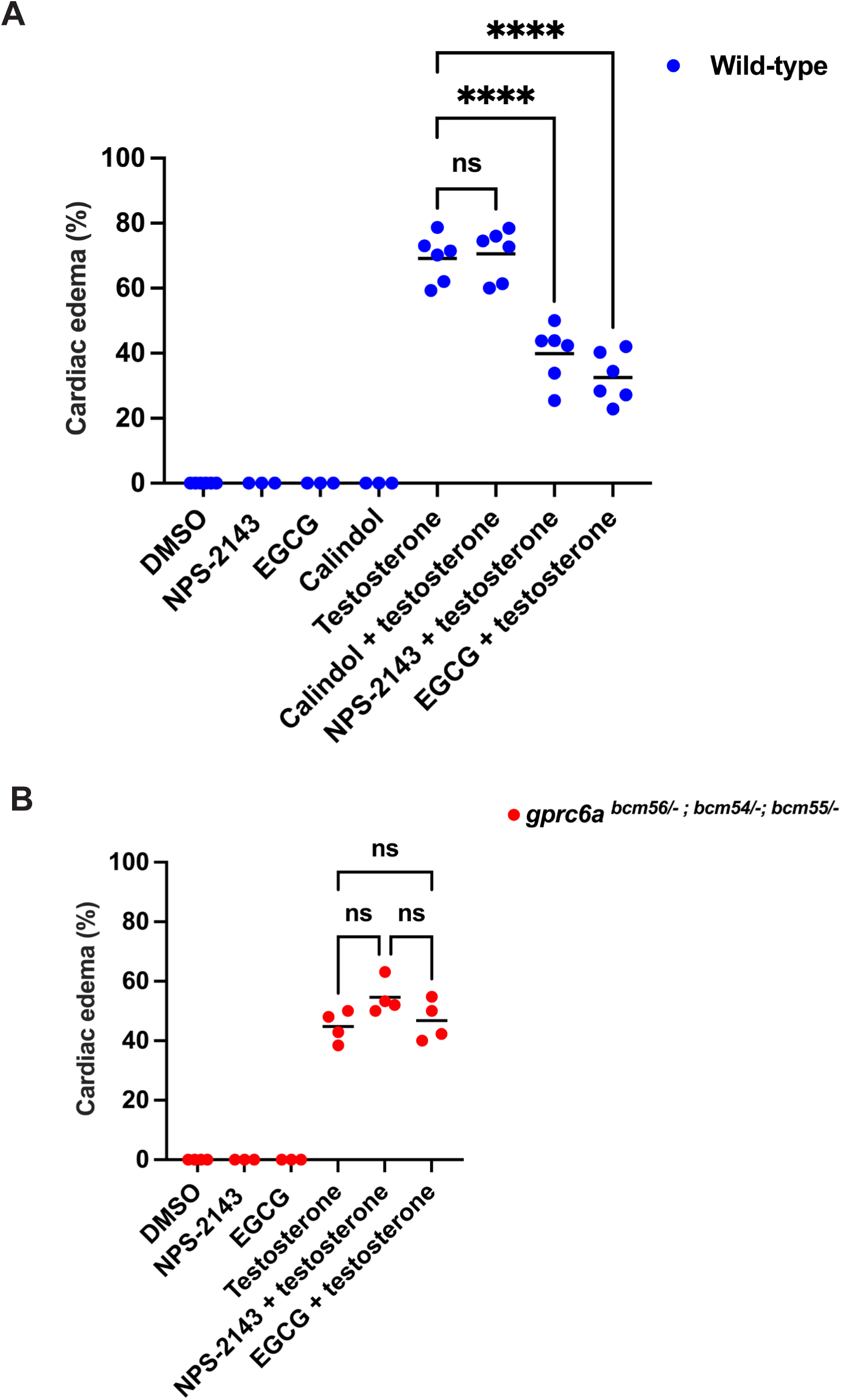
GPRC6A antagonists rescue testosterone phenotype in wild-type embryos but have no effect in *gprc6a* mutants. **(A**) Wild-type embryos were exposed to vehicle (0.1% DMSO) or GPRC6A antagonists: calindol (5 µM), NPS-2143 (5 µM), and epigallocatechin gallate (EGCG, 100 µM) with or without testosterone (30 µM) beginning at 3 hours post fertilization. Morphologic phenotypes were assayed at 3 days post fertilization. NPS-2143 and EGCG reduced cardiac edema following testosterone exposure. **(B**) Homozygous zygotic *gprc6a* mutant embryos were exposed to vehicle, NPS-2143 (5 µM), or EGCG (100 µM) with or without testosterone (30 µM) beginning at 3 hours post fertilization. Morphologic phenotypes were assayed at 3 days post fertilization. Antagonists failed to reduce cardiac edema in *gprc6a* mutant embryos. Each data point is the mean percent of embryos in a single clutch exhibiting cardiac edema (15-25 embryos per clutch). Clutches in the same treatment group were assayed on different days. Horizontal lines are the mean of each group. ****p < .0001, ns, not significant (p > .05), one-way ANOVA with Tukey’s multiple

**Table 4.**
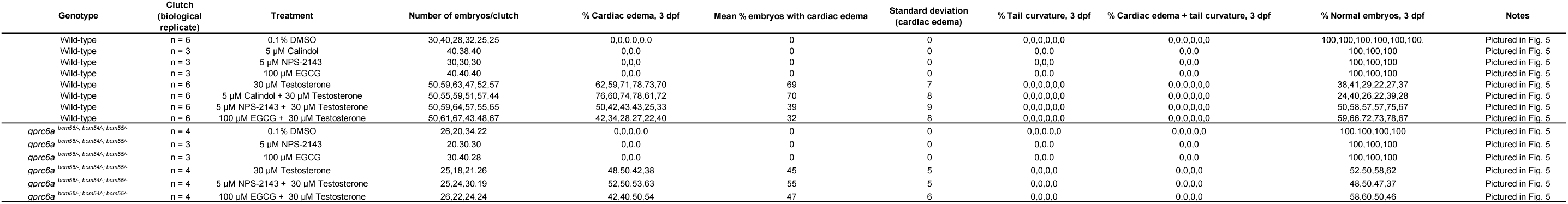
Embryo data accompanying Figure 7.

To test whether NPS-2143 and EGCG act specifically on zebrafish GPRC6A, we exposed homozygous *gprc6a* mutant embryos to testosterone alone or together with NPS-2143 and EGCG and assayed cardiac edema. There was no statistically significant difference in the percent of *gprc6a* homozygous embryos exhibiting cardiac edema following exposure to NPS-2143 or EGCG alone or in combination with testosterone (Fig. 7B, Table 4; testosterone mean embryos with cardiac edema 45% ± 5%, testosterone + NPS-2143 55% ± 5%, testosterone + EGCG 47% ± 6%, n=4 clutches, 18-30 embryos per clutch). These results show that the zebrafish *gprc6a* mutants do not respond to GPRC6A antagonists and supports the hypothesis that testosterone acts via GPRC6A to cause cardiac edema.

### Testosterone, and not a metabolite, causes cardiac edema in a GPRC6A-dependent manner

Testosterone is metabolized into molecules like estradiol. We showed that exposure to testosterone caused cardiac edema in a GPRC6A-dependent manner. However, it is not known whether testosterone, or a metabolite, is the active compound that causes the phenotype. We hypothesize that testosterone itself acts on GPRC6A to cause cardiac edema in zebrafish embryos. To test whether estradiol, a testosterone metabolite, causes a phenotype similar to testosterone, we exposed wild-type embryos at 3 hours post fertilization to 30 µM estradiol or 30 µM testosterone and assayed morphologic phenotypes at 3 days post fertilization. We found that exposure of wild-type embryos to estradiol did not cause cardiac edema or other morphologic phenotypes (Fig. 8A, Table 5), supporting the hypothesis that testosterone, and not estradiol, is the active compound that causes the phenotype.

**Figure 8.**
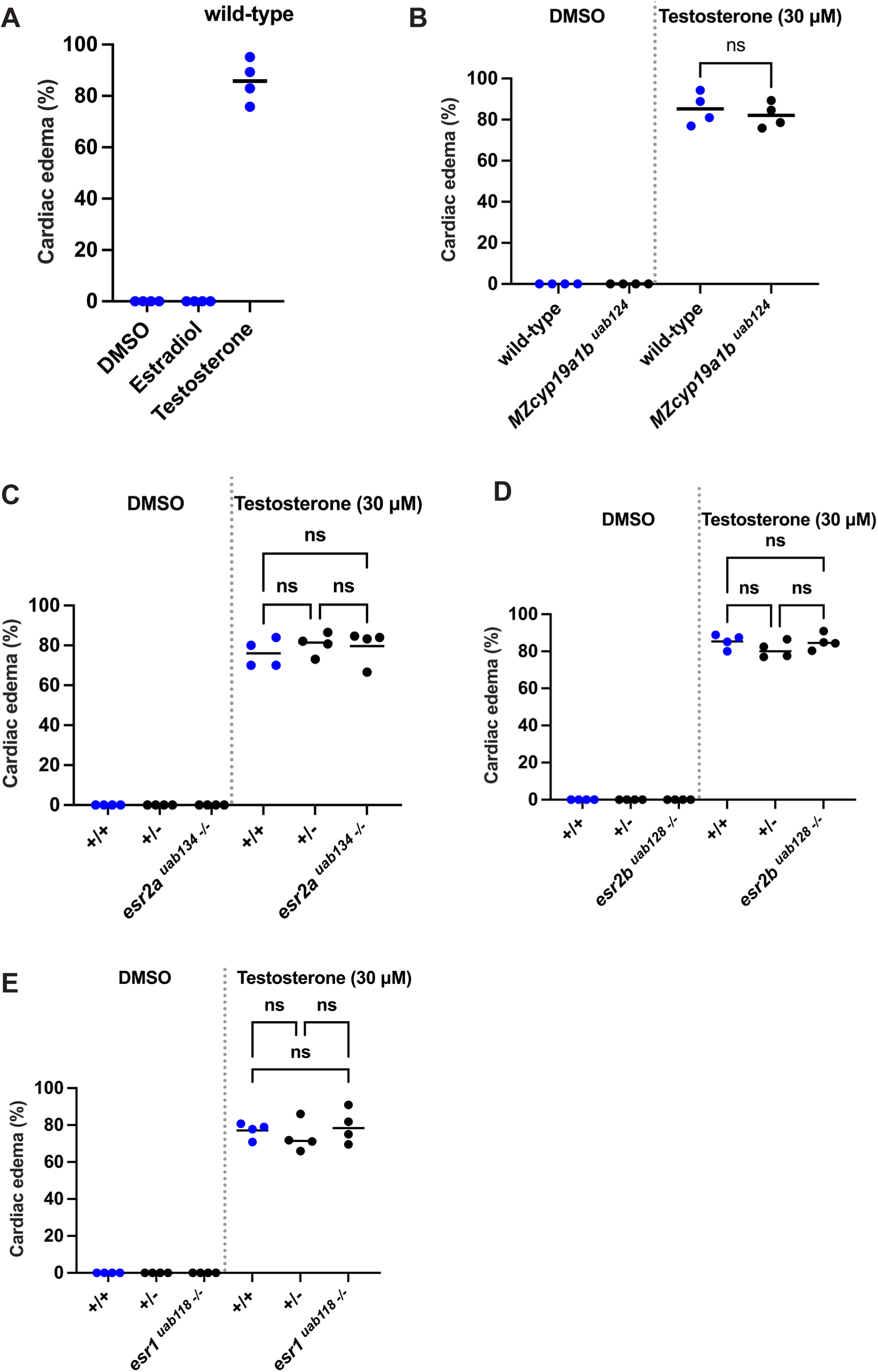
Testosterone directly acts on GPRC6A in ^E^ zebrafish embryos. **(A)** Wild-type embryos at 3 hours post fertilization were exposed to vehicle (0.1% DMSO), estradiol (30 µM) or testosterone (30 µM). Cardiac edema was assayed at 3 days post fertilization. **(B-E)** Zebrafish with mutations in aromatase (maternal zygotic *cyp19a1b*) or nuclear estrogen receptors (*esr2a*, *esr2b*, *esr1*) were exposed to vehicle or testosterone (30 µM) beginning at 3 hours post fertilization. Morphologic phenotypes were assayed at 3 days post fertilization. Each data point is the mean percent of embryos in a single clutch exhibiting cardiac edema (20-80 embryos per clutch). Clutches in the same treatment group were assayed on different days. Horizontal lines are the mean of each group. ns, not significant (p > .05), one-way ANOVA with Tukey’s multiple comparisons test.

**Table 5.**
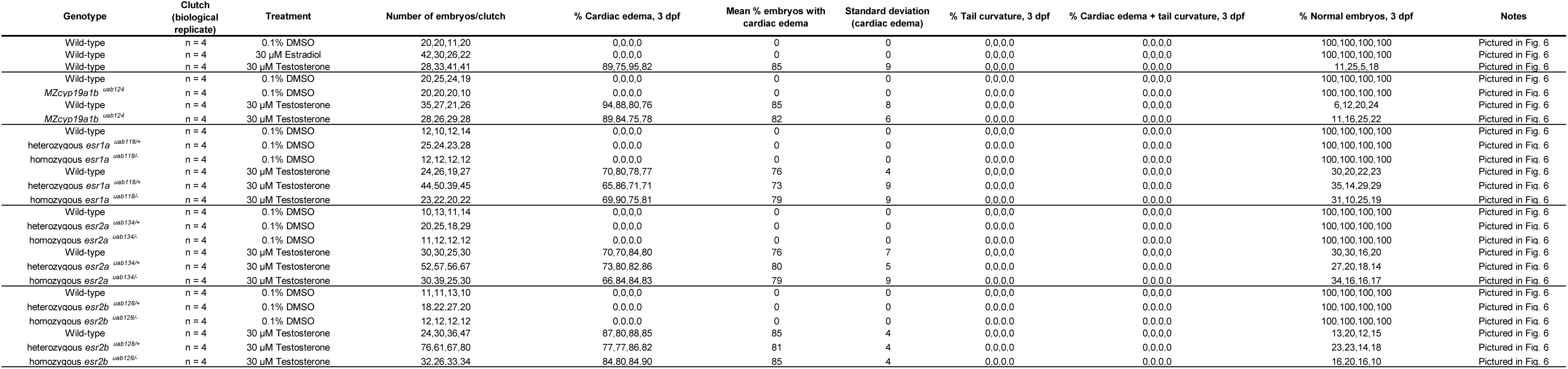
Embryo data accompanying Figure 8. Experiments within horizontal lines were performed together.

To further test whether testosterone acts on GPRC6A directly *in vivo*, we generated aromatase mutant embryos and asked whether testosterone causes a phenotype in an aromatase-dependent manner. Aromatase is the rate-limiting enzyme in the conversion of testosterone into estradiol (Conley and Hinshelwood, 2001; Duffy et al., 2010). Zebrafish have two aromatase genes, *cyp19a1a* and *cyp19a1b* (Chiang et al., 2001a; Chiang et al., 2001b). *cyp19a1a* is expressed exclusively in the gonads (which do not begin developing until ∼10 days post fertilization), while *cyp19a1b* is expressed in the brain at 2 days post fertilization (Chiang et al., 2001a; Chiang et al., 2001b; Lassiter and Linney, 2007). Here we focused on the *cyp19a1b* gene because of its expression during embryonic development.

Using CRISPR/Cas9, we generated heritable mutant alleles for *cyp19a1b* with a 20 bp insertion in exon 2 (Fig. S6A, maternal zygotic *cyp19a1b* mutant *MZcyp19a1b ^uab124^*). The mutation is predicted to result in a frameshift at amino acid 11 and a premature stop codon at amino acid 30 (Fig. S6B). We exposed *MZcyp19a1b ^uab124^* embryos to vehicle (0.1% DMSO) or testosterone at 3 hours post fertilization and assayed morphologic phenotypes at 3 days post fertilization. *MZcyp19a1b ^uab124^*embryos remained sensitive to testosterone exposure, similar to what we observed when wild-type embryos were exposed to testosterone (Fig. 8B, Table 5; wild-type mean embryos cardiac edema 85% ± 8%, *MZcyp19a1b* 82% ± 6%, n=4 clutches, 21-35 embryos per clutch). These results suggest that aromatase is not required for testosterone-dependent cardiac edema.

To exclude the possibility that testosterone acts via estrogen receptors (either via non-specific binding or via crosstalk between GPRC6A and nuclear estrogen receptors), we exposed estrogen receptor mutant embryos, *esr1a ^uab118/-^* (ERα), *esr2a ^uab134/-^* (ERβ1), and *esr2b ^uab128/-^* (ERβ2) to testosterone. We found no significant difference in the percent of embryos exhibiting cardiac edema phenotype between wild-type or heterozygous embryos versus homozygous *esr1a ^uab118/-^*, *esr2a ^uab134/-^*, and *esr2b ^uab128/-^* mutant embryos after testosterone exposure (Fig. 8C-E, Table 5). Taken together, these results suggest that testosterone, and not a metabolite, is the active agent that causes cardiac edema via GPRC6A.

### Genes regulated by T-GPRC6A

To investigate the mechanisms by which testosterone causes heart defects via GPRC6A, we performed RNA-sequencing (RNA-Seq) to identify genes regulated by T-GPRC6A. By comparing genes expressed in wild-type embryos following exposure to vehicle (0.1% DMSO), testosterone alone (30 µM), testosterone with EGCG (100 µM, GPRC6A antagonist) or testosterone with calindol (5 µM, calcimimetic with GPRC6A antagonism which was unable to rescue T-GPRC6A phenotype in Fig 7), we reasoned we could identify genes specifically regulated by T-GPRC6A and distinguish these from genes regulated by T-AR.

First, we generated 3 independent datasets from testosterone, calindol or EGCG treatment over DMSO (dataset 1: testosterone over DMSO; dataset 2: testosterone+calindol over DMSO; dataset 3: testosterone+EGCG over DMSO; Tables S3, S4, and S5). Each dataset was generated by defining a differentially expressed gene (DEG) as FDR < 0.05, irrespective of its fold change. We then made a list of T-GPRC6A specific DEGs by combining the significant genes that are differentially expressed in one direction (upregulated or downregulated in terms of fold change) in both testosterone and testosterone+calindol treated groups but were rescued in the testosterone +EGCG treated group by either showing no fold change or fold change in the opposite direction (e.g. if genes are upregulated in testosterone, they will be downregulated in testosterone+EGCG or vice versa, see Methods section for details). Using these approaches, we identified 655 T-GPRC6A-specific DEGs (Table S6).

To identify and prioritize candidate genes, we focused on genes that were downregulated by T-GPRC6A, reasoning that knockdown or mutation in these genes should mimic the testosterone exposure phenotype, while overexpression of these genes would rescue the testosterone exposure phenotype. We searched downregulated genes against the ZFIN database (Bradford et al., 2022) to identify genes that are expressed in the heart and genes for which a mutant or morpholino produced a cardiac phenotype similar to what we observe following testosterone exposure. We identified 37 genes expressed in the heart (Table S7). Among these, 7 exhibited a cardiac phenotype, of which 4 of them produced a phenotype similar to testosterone exposure when knocked down or mutated (Table S7).

### Overexpression of *grk3* or *pak1* mRNA partially rescues the testosterone exposure phenotype

We initially focused on candidate gene *grk3*. GRK3 has been predominantly recognized for its ability to desensitize GPCRs (Appleyard et al., 1999; Benovic et al., 1991; Boekhoff et al., 1994; Diverse-Pierluissi et al., 1996; Terman et al., 2004). This is achieved by phosphorylation of GPCR cytoplasmic domains and C termini that promote interactions between the phosphorylated GPCRs and *β*-arrestins (Benovic et al., 1987; Pitcher et al., 1992). This interaction sterically terminates G protein-mediated signaling. Zebrafish *grk3* morphants exhibited a cardiac phenotype similar to testosterone exposure (Jiang et al., 2009; Philipp et al., 2014). Thus, we would predict that downregulating or mutating *grk3* would increase the ability of GPRC6A to respond to testosterone, leading to cardiac abnormalities, while overexpressing *grk3* would reduce the ability of GPRC6A to respond to testosterone (presumably by desensitizing GPRC6A).

To test whether overexpression of *grk3* rescues the T-GPRC6A cardiac phenotype, we injected wild-type, 1-cell stage embryos with *grk3* mRNA and exposed them to testosterone or vehicle beginning at 3 hours post fertilization. We assayed morphology 3 days post fertilization. We found that embryos overexpressing *grk3* had a statistically significant reduction in cardiac edema following testosterone exposure compared to negative control embryos injected with *gfp* mRNA or uninjected embryos (Fig. 9; embryos injected with *grk3* mRNA mean cardiac edema 68% ± 4%, *gfp* mRNA mean cardiac edema 94% ± 1%, and uninjected embryos mean cardiac edema 93% ± 3%, n=3-5 clutches, 29-71 embryos per clutch). These rescue results support the hypothesis that testosterone acts via GPRC6A to cause cardiac phenotypes, while desensitizing GPRC6A via GRK3 overexpression prevents testosterone from exerting its deleterious effects on cardiac development (Fig. 10A).

**Figure 9.**
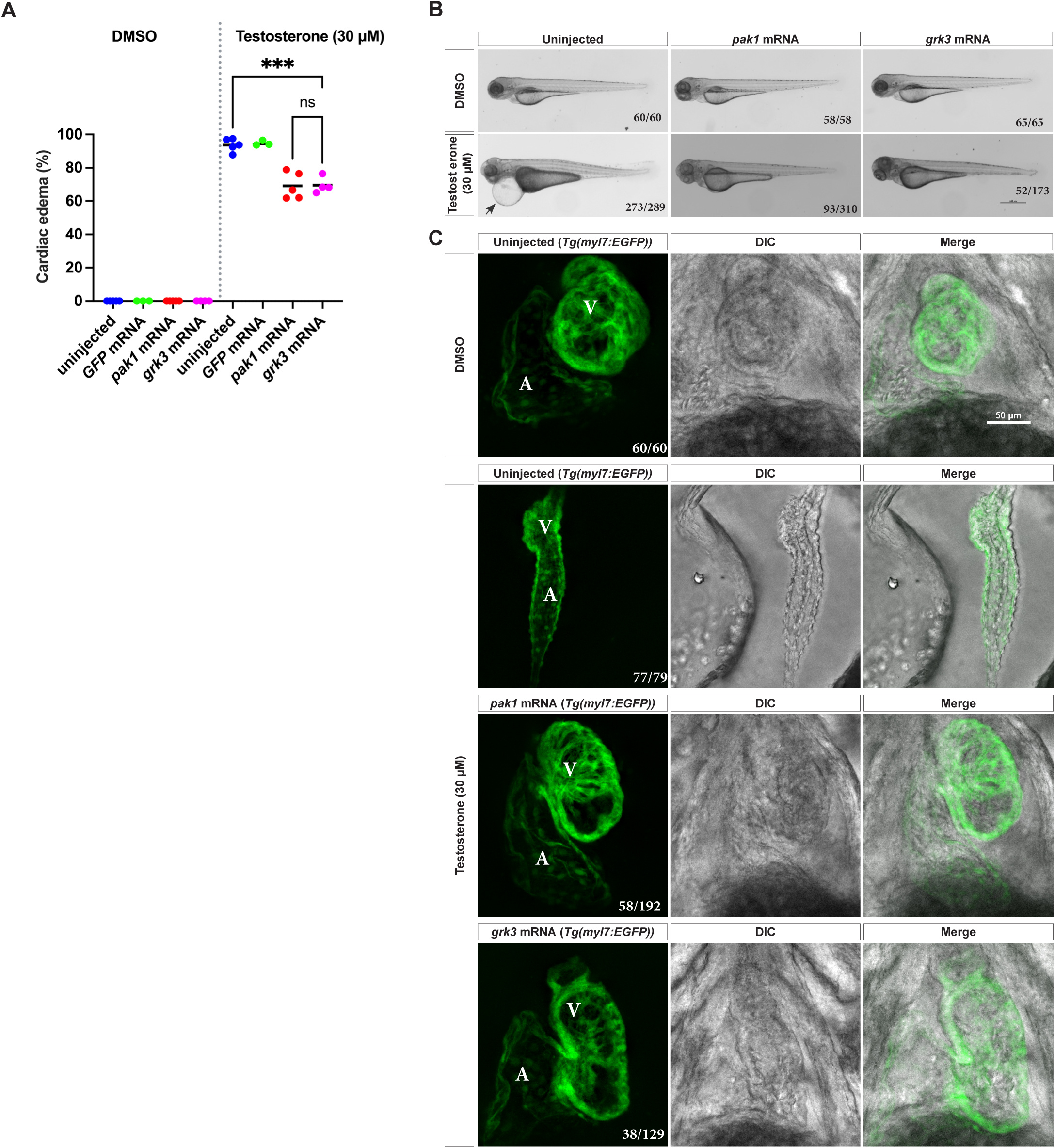
Testosterone phenotype is partially rescued by overexpression of *pak1* or *grk3* mRNA. **(A)** Wild-type embryos were injected with *gfp* mRNA, *pak1* mRNA, or *grk3* mRNA. Embryos were exposed to vehicle (0.1% DMSO) or testosterone (30 µM) beginning at 3 hours post fertilization (hpf). Morphologic phenotypes were assayed at 3 days post fertilization (dpf). *pak1* mRNA and *grk3* mRNA reduced cardiac edema following testosterone exposure, whereas *gfp* mRNA had no effect. Each data point is the mean percent of embryos in a single clutch exhibiting cardiac edema (29-71 embryos per clutch, 3-5 clutches per group). Clutches in the same treatment group were assayed on different days. Horizontal lines are the mean of each group. ****p < .0001, ns, not significant (p > .05), one-way ANOVA with Tukey’s multiple comparisons test. **(B)** Representative brightfield images of embryos from (A) at 3 days post fertilization. Note cardiac edema (arrows). Lateral views with anterior to the left, dorsal to the top. Fractions in the bottom right corners refer to the number of embryos with the indicated phenotype over the total number of embryos examined. Scale bar = 500 μm. **(C)** Cardiac phenotypes examined on the background of *Tg(myl7:EGFP)* embryos after exposure to testosterone (30 µM) or vehicle controls (0.1% DMSO) beginning at 3 hpf. Representative confocal maximum intensity projections of live embryo hearts at 3 dpf. Fractions in the bottom right corners refer to the number of embryos with the indicated phenotype over the total number of embryos examined (30-75 embryos per clutch, 2-3 clutches per group). Myocardial cells are labeled in green, A = atrium, V = ventricle, DIC = differential interference contrast, Scale bar = 50 μm.

**Figure 10.**
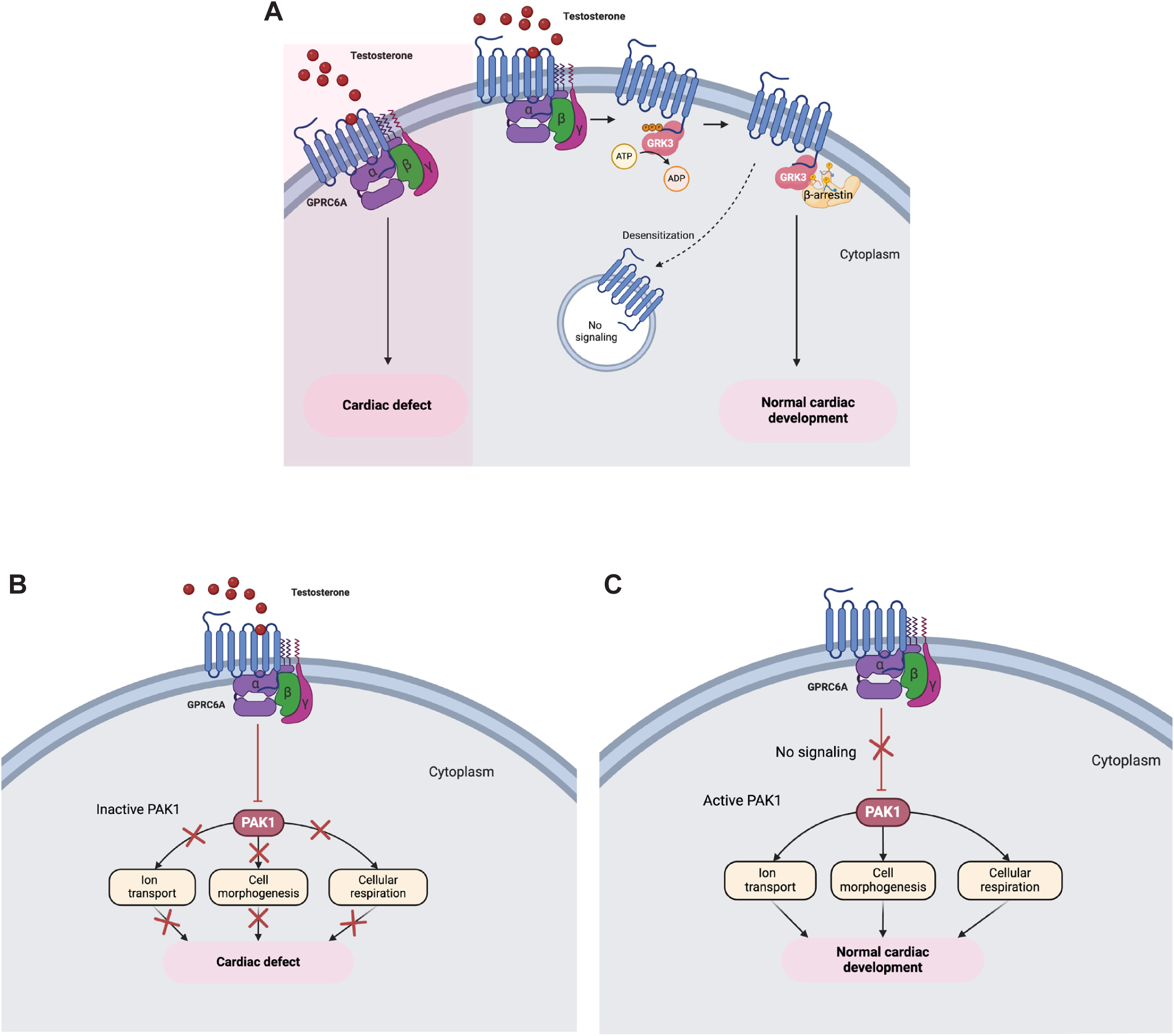
Model for GPRC6A-mediated signaling causing cardiac phenotype. **(A)** Testosterone activation of GPRC6A leads to activation of second messenger pathways that cause cardiac defects. The GPCR kinase GRK3 can reduce GPRC6A activity by promoting recruitment of β-arrestins, preventing G proteins from associating with GPRC6A, and increasing receptor internalization, blocking the effects of testosterone and restoring normal cardiac development. **(B)** Testosterone activation of GPRC6A inhibits or downregulates PAK1, thereby inactivating downstream signaling required for normal cardiac development. **(C)** In the absence of testosterone, GPRC6A is less active, so PAK1 is permitted to activate downstream signaling required for normal cardiac development.

We also focused on the candidate gene *pak1*. Zebrafish *pak1* knockdown embryos displayed a cardiac defect similar to testosterone-treated embryos (Jagadeeshan et al., 2017; Kelly et al., 2014; Lightcap et al., 2009). Whole-mount in situ hybridization revealed expression of *pak1* in the heart at 48 hpf (Kelly et al., 2014). *pak1* is a member of the serine/threonine kinase family, regulated by Ras-related GTPases RAC1 and CDC42 (Manser et al., 1994). PAK1 functions through multiple downstream proteins and signaling pathways, such as protein phosphatase 2A (Egom et al., 2016; MacDougall et al., 1991), ion channels (Egom et al., 2016; Wang et al., 2014), cardiac troponin I (Ke et al., 2004) and myosin light chain-2 (Monasky et al., 2012). In our RNA-seq data, not only was *pak1* downregulated by T-GPRC6A (Table S6), but also several ion channels involved in zebrafish cardiac function that are plausible PAK1 targets, including *scn5Laa, itpr1b, kcnn3, kcnj14*, and *ftr01*, were downregulated by T-GPRC6A (Table S6 and S7). Furthermore, a gene set enrichment analysis (GSEA) revealed several ion transport/signaling pathways, consistent with PAK1 function, that were downregulated in both the testosterone and testosterone+calindol treated groups and rescued in the testosterone+EGCG treated group (Table S8, sodium ion transmembrane transport, calcium-mediated signaling, calcium ion transmembrane transport, cation transmembrane transport, among others). These findings encouraged us to prioritize *pak1* as a candidate.

To test whether overexpression of *pak1* rescues the T-GPRC6A cardiac phenotype, we injected wild-type, 1-cell stage embryos with *pak1* mRNA and exposed them to testosterone or vehicle beginning at 3 hours post fertilization. We assayed morphology 3 days post fertilization. Similar to overexpressing *grk3*, we found that embryos overexpressing *pak1* also had a statistically significant reduction in cardiac edema following testosterone exposure compared to negative control embryos injected with *gfp* mRNA or uninjected embryos (Fig. 9; embryos injected with *pak1* mRNA mean cardiac edema 66% ± 8%, *GFP* mRNA mean cardiac edema 94% ± 1%, and uninjected embryos mean cardiac edema 93% ± 3%, n=3-5 clutches, 29-71 embryos per clutch). We conclude that GPRC6A, when activated by testosterone, inhibits PAK1 activity, leading to abnormal cardiac development (Fig. 10B, C).

## Discussion

We demonstrate that a CRISPR screen in zebrafish embryos can be used to identify genes required for testosterone-dependent phenotypes. The zebrafish screening approach complements existing cell-based approaches. Membrane steroid receptors are typically identified based on their binding or activity in cultured cells, and then tested for similar activity *in vivo*. It can be difficult to determine whether a receptor responds to steroids *in vivo*, especially receptors that have multiple ligands. For example, the estrogen receptor modulator tamoxifen was shown to bind directly to the human sodium channel Nav1.2 and inhibit sodium conductance *in vitro* (Sula et al., 2021), but whether this occurs *in vivo* is not known. The zebrafish screening approach allows for the direct identification of membrane steroid receptors *in vivo*.

Our approach relies on using CRISPR-Cas9 to rapidly generate mutant zebrafish. To generate heritable mutants, germline transmitting animals need to be identified and mated to produce homozygous embryos, a process that takes months. In contrast, analyzing gene function in mosaic mutant animals (F0 generation) takes less than a week. Our approach was enabled by recent findings that injecting zebrafish embryos with Cas9 protein and a cocktail of 3-4 guide RNAs targeting the same gene induces mutations at a high enough burden to observe phenotypes in mosaic mutant embryos (Klatt Shaw and Mokalled, 2021; Kroll et al., 2021; Wu et al., 2018). One potential limitation of CRISPR screens is the failure of F0 mosaic mutants to recapitulate phenotypes found in heritable, non-mosaic mutants (eg F1 and subsequent generations). In our screen, this was not the case. We identified similar phenotypes in F0 mosaic and in non-mosaic *gprc6a* mutants (compare Figures 4 and 6), demonstrating the validity of using the F0 CRISPR screening approach.

Our pilot screen tested 4 candidate genes using CRISPR-Cas9. However, new barcoding and microfluidics approaches, such as MIC-Drop, could allow us to screen and mutate hundreds or thousands of genes (Parvez et al., 2021). Combining different chemical exposures with MIC-Drop offers the unprecedented ability to comprehensively identify membrane receptors for any steroid molecule that causes a phenotype in zebrafish embryos. Here, we assayed gross, easily observable morphologic phenotypes. It is also possible to assay more subtle phenotypes, for example by using transgenic reporter zebrafish that mark a cell-type of interest, or by using assays to measure behaviors like locomotor activity.

We found that testosterone exposure caused cardiac edema and heart malformations in approximately 80% of wild-type embryos. This phenotype was partially rescued by co-administration of two different GPRC6A antagonists (Figure 7). The testosterone phenotype was also partially rescued in *gprc6a* mutants (both mosaic F0 and heritable homozygotes, Figure 4, Figure 6). Similar results using F0 mosaic mutants, F0 non-mosaic mutants, and pharmacologic antagonists, plus rescue with overexpression of *grk3* and *pak1* (Figure 9), suggests the partial rescue is not an artifact of mutagenesis. What explains the lack of a complete rescue? Testosterone could also be acting simultaneously through multiple pathways, for example by binding to additional receptors or by altering the fluidity of cell membranes or the mobility of membrane proteins (Liang et al., 2001; Shivaji and Jagannadham, 1992; Whiting et al., 1995; Whiting et al., 2000). The fact that the testosterone phenotype in *gprc6a;ar* double mutants was similar to *gprc6a* single mutants (Figure 5) suggests that T-GPRC6A acts independently of AR in this case.

Our results are the first to identify a role for androgen-dependent GPRC6A activity during embryonic development. Our results are consistent with results from cultured cells and mouse models demonstrating that GPRC6A responds to testosterone. In HEK-293 cells, testosterone stimulated extracellular signal-regulated kinase (ERK) activity, which was blocked by GPRC6A antagonists (Pi et al., 2018). In wild-type mice, testosterone treatment increased levels of phosphorylated ERK and Egr-1 expression in bone marrow and testis, which was reduced or absent in GPRC6A mutant mice (Pi et al., 2010). Several ligands activate GPRC6A, including calcium, magnesium, and the peptide hormone osteocalcin (Pi et al., 2005). It remains to be seen whether or how testosterone influences the response of GPRC6A to other ligands. Osteocalcin is maternally deposited into zebrafish oocytes and expressed at 2 days post fertilization (Bensimon-Brito et al., 2012), developmental stages when embryos exposed to testosterone exhibit cardiac phenotypes. Future studies could explore whether osteocalcin is required for testosterone-GPRC6A-dependent cardiac phenotypes.

We also discovered part of the mechanism by which T-GPRC6A regulates cardiac development, via the serine/threonine kinase PAK1. Overexpression of *pak1* partially rescued the testosterone exposure phenotype in wild-type embryos (Fig 9). A previous report found that zebrafish pak1 signals through the Erk pathway and the transcription factor Gata6 to promote proper heart development (Kelly et al., 2014). Our results are consistent with this observation. We propose that exogenous testosterone activates GPRC6A, which then inhibits Pak1 and leads to abnormal heart development (Fig. 10). In the absence of testosterone, Pak1 is free to activate its downstream targets, which may include Erk/Gata6 but could also include ion channels and transporters required for normal heart development. Using RNA-seq, we identified several ion channel genes downregulated by T-GPRC6A. While the interaction between these ion channels and *pak1* has not been investigated in zebrafish, they could play a role in cardiac development and function similar to ion channels known to function downstream of Pak1 in other organisms.

Our ELISA results show that an average of ∼60 nM testosterone per embryo causes a phenotype. We failed to detect any testosterone in untreated embryos. We hypothesize that under normal conditions, zebrafish embryos at 0-3 dpf contain low levels of testosterone, which allow the heart to develop normally (Fig 10). Increasing testosterone levels activates GPRC6A, which leads to improper heart development. Because the zebrafish gonad does not differentiate until approximately 10 dpf, it is likely that endogenous levels of testosterone are low until that time. It is possible that in zebrafish, the heart (and other organs) are more sensitive to testosterone exposure during early periods of differentiation and development. Once developed and more mature, the heart may be less susceptible to potentially deleterious effects of testosterone exposure. This idea of a critical window of sensitivity is consistent with data from sheep. Pregnant ewes exposed to increased testosterone between gestational days 30 and 59 produced fetuses with smaller hearts due to reduced cardiomyocyte maturation and proliferation (fetuses were delivered at 135 days). In contrast, pregnant ewes exposed to increased testosterone between gestation days 60 and 90 had normal cardiac growth and development (Jonker et al., 2018).

Gestational testosterone excess, such as occurs in polycystic ovarian syndrome (Sir-Petermann et al., 2002), is associated with fetal abnormalities that increase risks for heart dysfunction later in life. Fetal sheep exposed to increased testosterone exhibited myocardial disarray in the left ventricle, increased cardiomyocyte size, increased expression of genes associated with cardiac hypertrophy, and fetal growth retardation (Ghnenis et al., 2022; Manikkam et al., 2004; Ramamoorthi Elangovan et al., 2023; Vyas et al., 2016). In rats, maternal testosterone excess was associated with cardiac hypertrophy in offspring (Hou et al., 2019). While the molecular mechanisms of how testosterone causes these phenotypes is not well understood, the assumption is that these phenotypes are caused by testosterone activating canonical nuclear androgen receptors. Our results in zebrafish suggest that gestational testosterone excess may also cause deleterious effects via membrane androgen receptors like GPRC6A. Furthermore, studying the effects of excess androgens in zebrafish embryos could provide insights into how gestational hyperandrogenism associated with polycystic ovarian syndrome affects fetal development.

Our chemical-genetic F0 CRISPR screening approach has several limitations. Our scheme relies on identifying phenotypes following exposure to steroids, and then testing whether a gene mutation rescues the exposure phenotype (Figure 1). If a mutant has a phenotype in the absence of chemical exposure, then we cannot test that gene. For example, embryos injected with gRNAs targeting *cav1.2* exhibited cardiac edema in the absence of chemical exposure. Consequently, we could not test whether *cav1.2* mutations rescue testosterone-dependent cardiac edema. This also demonstrates the strength of the approach. The phenotype we observed in *cav1.2* mosaic mutants was similar to the phenotypes observed in *cav1.2* heritable, homozygous mutants (Rottbauer et al., 2001), further supporting the idea that F0 mosaic mutants recapitulate phenotypes observed in heritable, non-mosaic mutants.

Another limitation is that genetic compensation could produce false negative results. A mutation in a gene could lead to upregulation of a related gene that compensates for the original mutation. In this case, we would falsely conclude that the target gene is not involved in steroid signaling. Future studies can overcome this limitation by performing a deletion screen, using two gRNAs flanking the target gene to delete the entire target gene. This would prevent production of mRNA and consequently would reduce the occurrence of compensation triggered by mutant mRNA (El-Brolosy et al., 2019; Fernandez-Abascal et al., 2022; Ma et al., 2019; Serobyan et al., 2020). Another concern are phenotypes resulting from the off-target effects of Cas9. These can be validated by performing RNA rescue studies or by generating heritable, non-mosaic mutations in the target gene. By successively breeding such mutants with wild-type animals, one selects for the target mutation and loses potential off-target mutations. This is the strategy we employed. We found that homozygous *gprc6a* mutants are partially resistant to testosterone exposure (Figure 6), similar to what we observed in mosaic *gprc6a* F0 mutants (Figure 4). Thus, the off-target effects of Cas9 are unlikely to influence our results.

In conclusion, we developed a combined chemical and genetic screening approach to identify genes required for androgen-dependent phenotypes in zebrafish embryos. Our pilot screen revealed that testosterone acts via GPRC6A to cause cardiac phenotypes. This chemical-genetic screening platform can be adapted to discover genes encoding membrane proteins that act as receptors for any steroid or related small molecule.

## Acknowledgements

We thank Lauren Pandolfo and the staff of the BCM zebrafish facility for caring for our zebrafish, the members of NR IMPACT for scientific feedback during the early phases of this project, and members of the Gorelick laboratory for advice on experiments. This work was supported by the NIH Training Program in Precision Environmental Health Sciences T32 ES027801. The Baylor College of Medicine high-performance computing cluster, managed by the Biostatistics and Informatics Shared Resource, is supported by grants from NCI P30-CA125123 and Institutional funds from the Dan L Duncan Comprehensive Cancer Center and Baylor College of Medicine. Figures 1 and 10 were created using Biorender.com.

---

**Figure S1.**
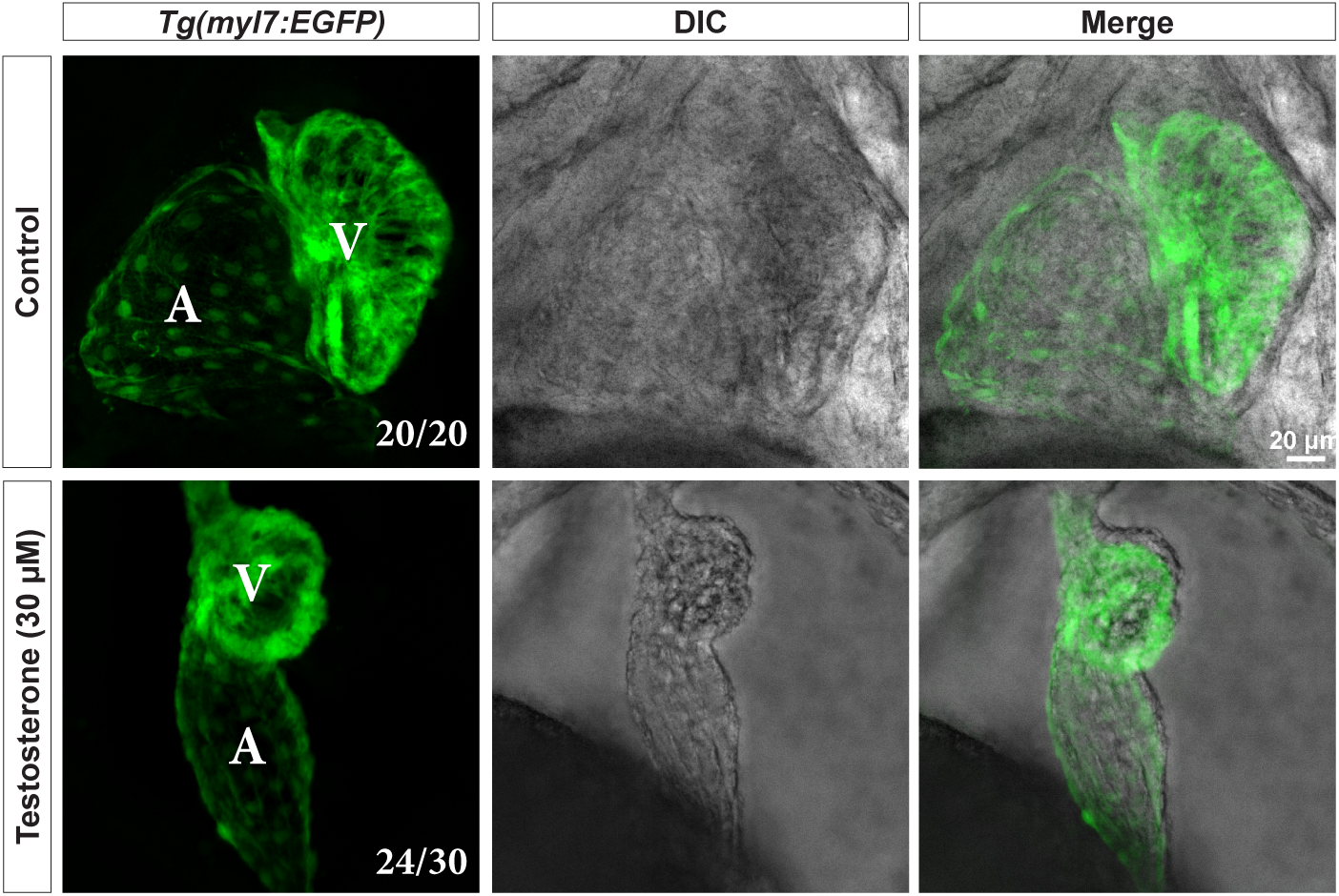
Testosterone exposure causes elongated atrium, shrunken ventricle and failure of heart looping. Confocal maximum intensity projections of live *Tg(myl7:EGFP)* embryos focused on the heart at 3 days post fertilization. Embryos were exposed to testosterone (30 μM) or vehicle control (0.1% DMSO) beginning at 3 hours post fertilization. 100% of control embryos had normal hearts (n=20), 80% of embryos exposed to testosterone had abnormal hearts (24 out of 30). Myocardial cells are labeled in green, A = atrium, V = ventricle, DIC = differential interference contrast.

**Figure S2.**
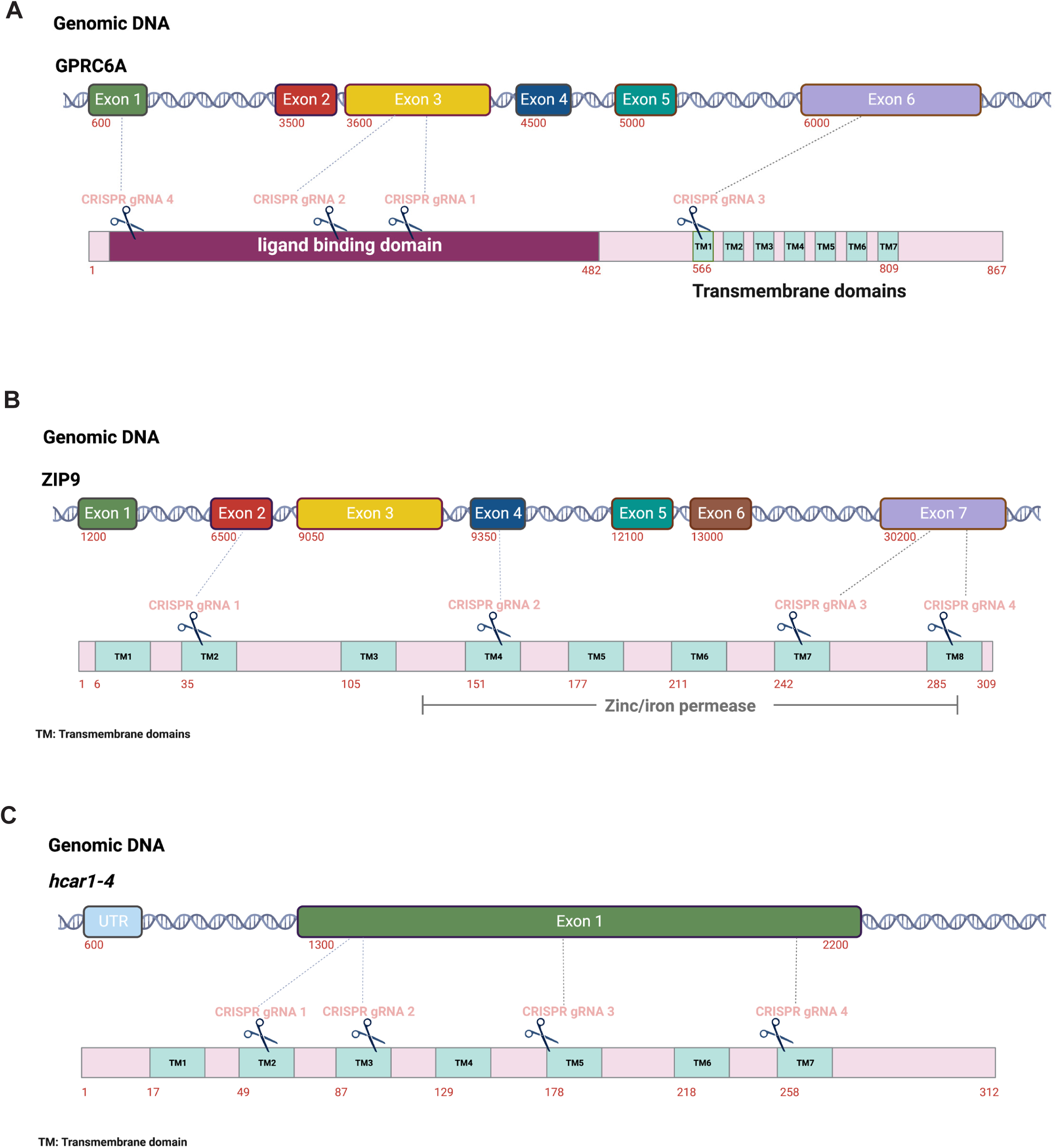
Location of guide RNAs targeting putative membrane androgen receptor genes. Four guide RNAs each were used to target *gprc6a* (**A**), *zip9* (**B**), and *hcar1-4* (**C**) genes. The predicted location in the corresponding protein is shown below each genomic DNA. Numbers indicate the starting and ending nucleotides of exons in the genomic DNA or amino acid numbers in the protein sequence. Boxes in the protein sequence indicate key structural or functional domains. Figure created using Biorender.com.

**Figure S3.**
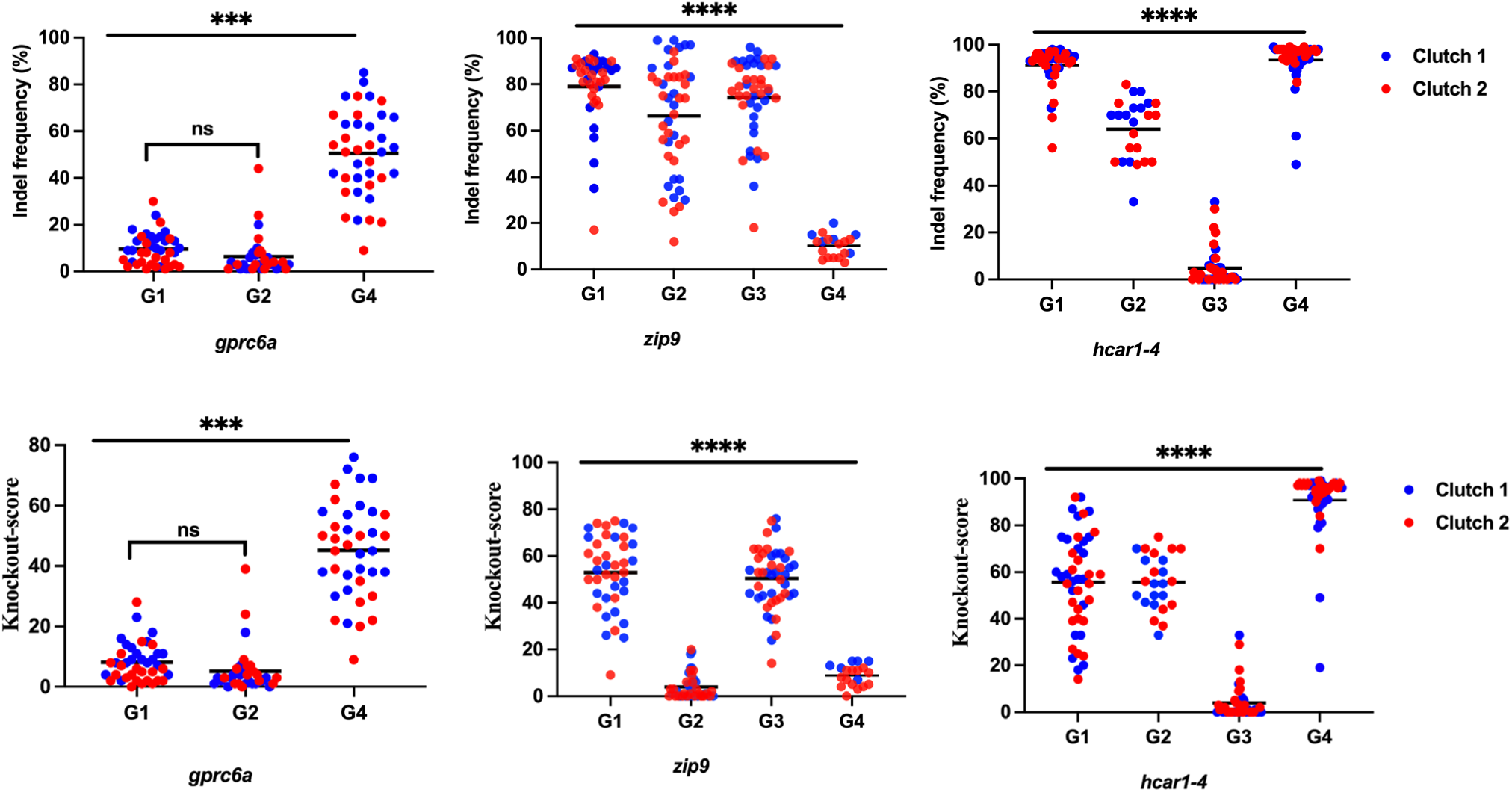
Multiplexed gRNA injection results in frequent mutations in injected embryos. 1-cell stage zebrafish embryos were injected with Cas9 protein together with a cocktail of 4 guide RNAs (gRNA 1 = G1; gRNA 2 = G2; gRNA 3 = G3; gRNA 4 = G4) targeting different regions of the indicated genes (*gprc6a, zip9, hcar-1-4*). At 1 day post fertilization, we harvested genomic DNA from each embryo individually, performed PCR to amplify each of the 4 different regions targeted by each guide RNA, and then we Sanger sequenced each PCR product. Sanger sequence was deconvoluted and analyzed using the ICE (Inference of CRISPR Edits) algorithm. Top graphs, percent of sequence from each embryo that has a mutation (indel frequency). Bottom graphs, percent of mutations in sequencing reactions from each embryo predicted to produce a non-functional protein (knockout score). Each data point represents sequencing results from a single embryo. Data points of the same color represent embryos from the same clutch, embryos that were produced from the same parents and injected and assayed together on the same day. Horizontal lines are the mean for each group. G3 targeting *gprc6a* produced no mutations and is not shown. ***p < .001, ****p < .0001, ns, not significant (p > .05), one-way ANOVA with Tukey’s multiple comparisons test.

**Figure S4.**
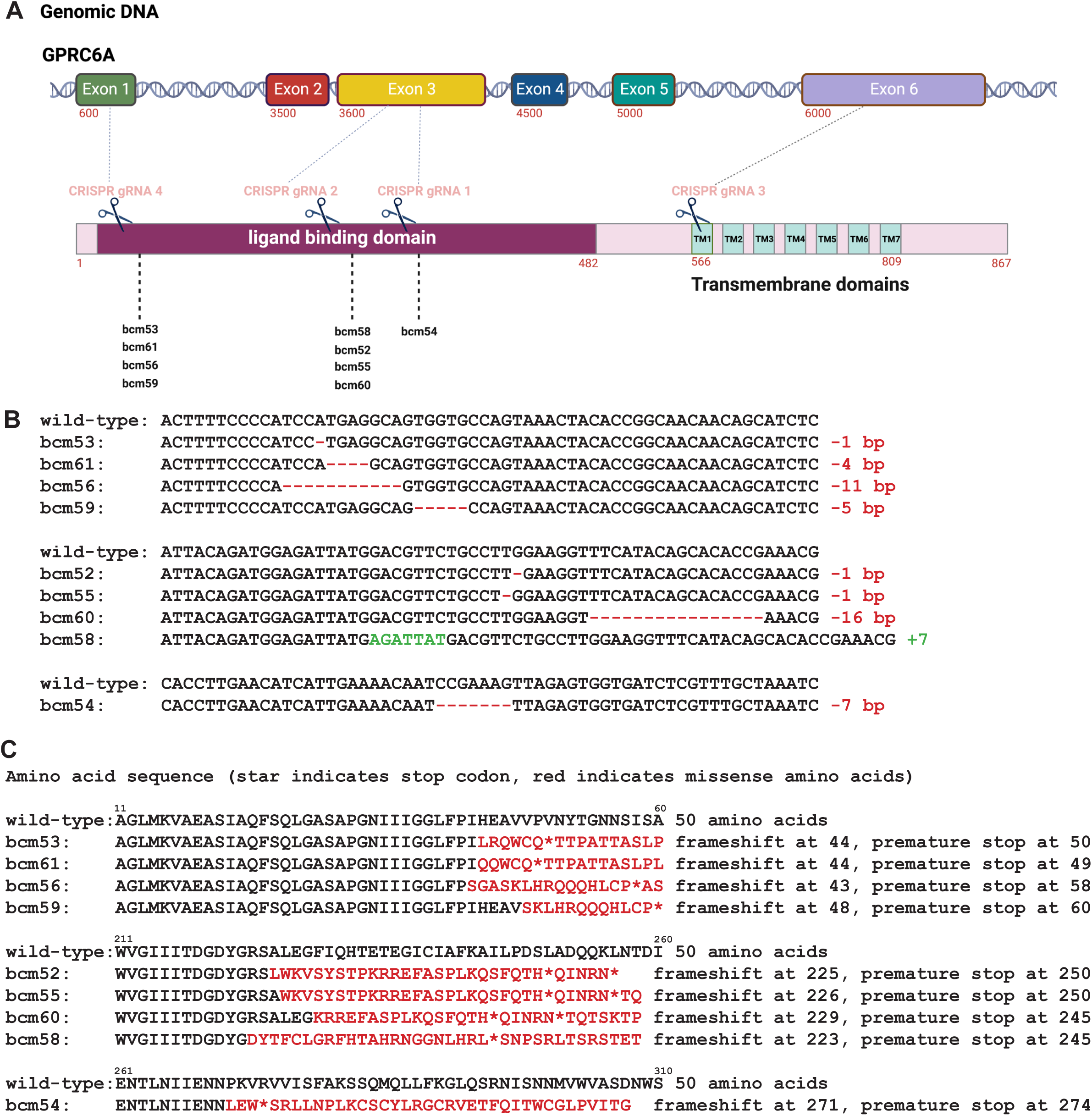
Generation of heritable *gprc6a* mutant zebrafish. **(A)** We generated heritable mutant alleles for *gprc6a* using CRISPR/Cas9. **(A)** 1-cell stage embryos were injected with 4 gRNAs targeting sites within exons 1, 3 and 6 of *gprc6a* (gray dotted lines). Injected embryos were raised to adulthood and crossed to wild-type zebrafish. Resulting progeny were raised to adulthood. We used PCR and Sanger sequencing to identify heterozygotes with putative loss-of-function mutations in the predicted protein, illustrated below the genomic DNA. Zebrafish containing desirable mutations were given allele designations (*bcm53*, *bcm61*, *bcm56*, *bcm59*, *bcm58*, *bcm52*, *bcm55*, *bcm60*, *bcm54*) and propagated. Numbers indicate the starting and ending nucleotides of exons in the genomic DNA or amino acid numbers in the protein sequence. Boxes in the protein sequence indicate key structural or functional domains. **(B)** Nucleotide sequence for each allele was confirmed by Sanger sequencing. Red dashes indicate deletions, green nucleotides indicate insertions. (**C)** Predicted changes to

**Figure S5.**
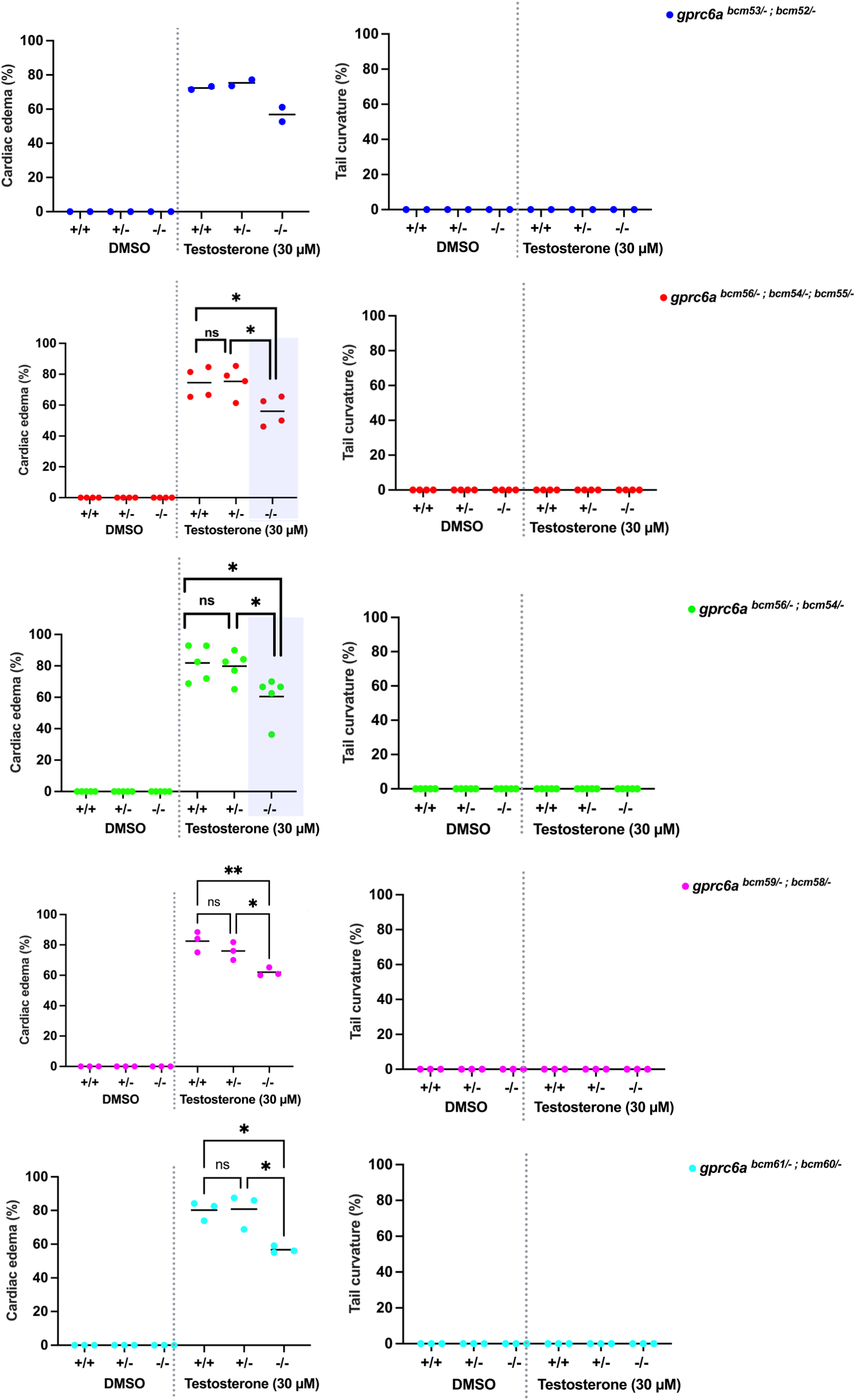
Testosterone phenotypes in homozygous *gprc6a* mutant embryos based on allele. Percent of embryos exhibiting cardiac edema (left graphs) or tail curvature (right graphs) phenotypes. Each data point is the mean percent of embryos in a single clutch exhibiting cardiac edema or tail curvature (15-50 embryos per clutch). Clutches in the same treatment group were assayed on different days. Horizontal lines are the mean of each group. This is the same data from Figure 6B but each allele is graphed separately. Genotypes and treatments are separated by vertical gray dotted lines in each graph. * p < .05, ** p < .01, ns,

**Figure S6.**
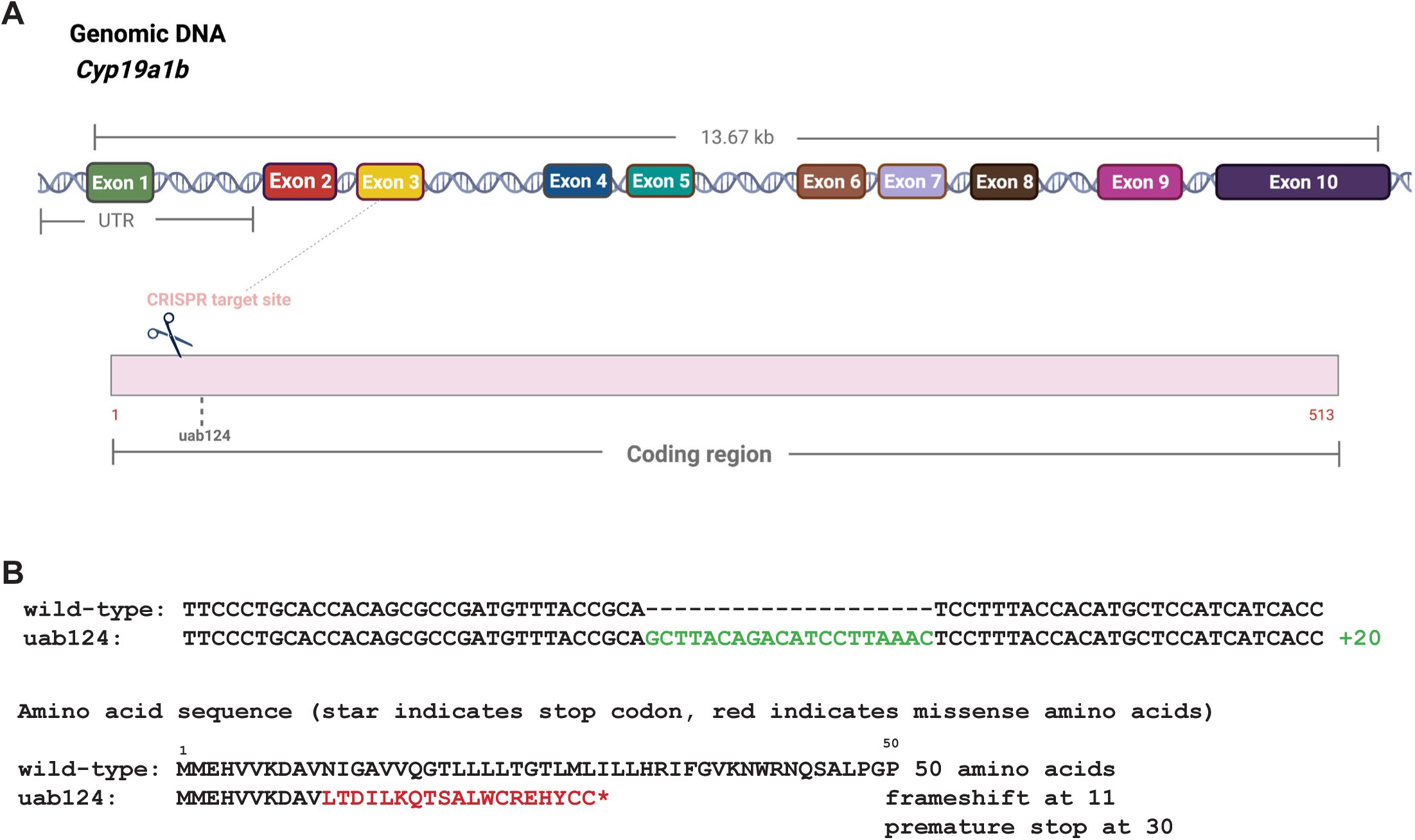
Generation of *cyp19a1b* mutant zebrafish. We generated heritable mutant alleles for *cyp19a1b* using CRISPR/Cas9. **(A)** 1-cell stage embryos were injected with gRNA targeting exon 3 of *cyp19a1b* (gray dotted lines). Injected embryos were raised to adulthood and crossed to wild-type zebrafish. Resulting progeny were raised to adulthood, where we identified allele *uab124*. Numbers indicate the starting and ending amino acid numbers in the protein sequence. **(B)** *uab124* nucleotide sequence was confirmed by Sanger sequencing. Green nucleotides indicate insertions. Predicted changes to amino acid sequence are also indicated. Red indicates amino acid mutations, and asterisk indicates premature stop.

## Notes

### Competing Interest Statement

Daniel A Gorelick is the editor in chief of the journal Biology Open. The other authors have no competing interests to declare.

### Summary of Updates

New RNA-seq data (supplemental files updated, Figure 9), section on testosterone uptake updated to include ELISA results (Figure 3)

